# Enabling large-scale genome editing by reducing DNA nicking

**DOI:** 10.1101/574020

**Authors:** Cory J. Smith, Oscar Castanon, Khaled Said, Verena Volf, Parastoo Khoshakhlagh, Amanda Hornick, Raphael Ferreira, Chun-Ting Wu, Marc Güell, Shilpa Garg, Hannu Myllykallio, George M. Church

## Abstract

To extend the frontier of genome editing and enable the radical redesign of mammalian genomes, we developed a set of dead-Cas9 base editor (dBE) variants that allow editing at tens of thousands of loci per cell by overcoming the cell death associated with DNA double-strand breaks (DSBs) and single-strand breaks (SSBs). We used a set of gRNAs targeting repetitive elements – ranging in target copy number from about 31 to 124,000 per cell. dBEs enabled survival after large-scale base editing, allowing targeted mutations at up to ~13,200 and ~2610 loci in 293T and human induced pluripotent stem cells (hiPSCs), respectively, three orders of magnitude greater than previously recorded. These dBEs can overcome current on-target mutation and toxicity barriers that prevent cell survival after large-scale genome engineering.

**One Sentence Summary:** Base editing with reduced DNA nicking allows for the simultaneous editing of >10,000 loci in human cells.

---

Endogenous transposable elements (TEs) such as Alu(*1*), Long Interspersed Elements-1 (LINE-1)(*2*–*4*) or Human Endogenous Retro Viruses (HERV)(*5*) make up ~45% of the human genome(*6*). While originally characterized as “junk DNA,” TEs are now recognized as having shaped the evolution of the human genome, and their residual transposition activity has been linked to human physiology and disease. For instance, LINE-1 sequences (17% of the genome) are highly active in neurons(*7*), can disrupt gene expression(*4*), and are suspected of having roles in human neurological diseases(*2*, *3*, *8*, *9*) and aging(*10*, *11*). Alu and HERV have been associated with aging(*12*) and multiple sclerosis(*5*, *13*) respectively. The most direct test of such hypotheses would involve genomically inactivating these elements, but this has been effectively out of reach because it would require editing large numbers of distinct loci, challenging the capacity of current editing methods and the ability of cells to tolerate their activity due to the high toxicity of DSBs(*14*, *15*). The current record for simultaneous inactivation of TEs – 62 elements – was achieved using CRISPR/Cas9(*16*) on Porcine Endogenous Retroviruses (PERVs) in a transformed pig cell line. Two years later a live pig was born with genome-wide KO of all 25 PERVs(*17*).

CRISPR/Cas9 incurs toxicity because it generates double-strand DNA breaks (DSBs)(*14*). These DSBs contribute to its high genome-editing efficiency by potently triggering endogenous processes that repair them with random or user-specified variations, but high numbers of concurrent DSBs overwhelm these processes and cause cell death. Recently, however, two types of CRISPR/Cas9 “base editors” (BEs) were developed (*table S1*) by fusing variants of Cas9 that are either “dead” (dCas9; both nuclease domains inactivated) or “nicking” (nCas9; one nuclease domain inactivated), in which the DSB-generating nuclease domains are disabled, to a nucleotide deaminase. Cytidine base editors (CBEs: either dCBEs or nCBEs(*18*)) employ cytidine deaminases and convert C:G base pairs to T:A, while adenine base editors (ABEs: either dABEs or nABEs(*19*)) use adenine deaminases and convert A:T base pairs to G:C. Using properly designed gRNAs, C->T conversions may be used to generate stop codons to knock-out protein coding genes of interest(*14*). Improvements in base editing purity – the frequency of desired base conversion within a target window – have been achieved by fusing bacterial mu-gam protein to the base editor to generate nCBE4-gam(*20*). Naming conventions for all BEs are summarized in *table S2*.

To achieve similar efficiencies to native Cas9 all base editor generations beyond the first are nBEs. As a result, base editing has been broadly demonstrated with high efficiency in a range of species including human zygotes(*21*). A main motivation for developing BEs that avoid DSBs was to reduce the level of random vs. user-specified mutations caused by “live” Cas9, but the reduced toxicity of BEs accrued by avoiding DSBs has also facilitated the editing of single targets in sensitive cell types such as hiPSCs(*22*). However, whether these BEs can enable concurrent editing in human cells of sites as numerous as high copy TEs has not been explored but is particularly relevant to genome wide recoding efforts such as genome project write(*23*) (GP-write). To recode the human genome would require an estimated 4438 to 9811 precise modifications to remove all instances of one of the three stop codons(*24*).

## gRNA design and copy number estimation of transposable elements

To assess the efficiency and toxicity of current editing technologies as applied to TEs, we designed and tested gRNAs against Alu, LINE-1, and HERV which vary in copy number from 30 to greater than 100,000 across the genome (Fig. 1A). Alu and LINE-1 gRNAs were respectively designed on the consensus sequences obtained from repeatmasker(*25*) (*table S3*) and on the consensus of 146 full-length sequences that encodes both functional ORF1 and ORF2 proteins(*26*). Finally, gRNAs against HERV-W, one subfamily of HERV, were designed on the consensus of putatively active retro-viruses(*27*) (*table S3)*. We performed qPCRs of genomic DNA (gDNA) generated using consensus sequence-based primers to estimate the relative abundance in HEK 293T and PGP1 iPSCs (Fig 1A). The copy number of HERV-W, LINE-1, and Alu elements at the edited sites were respectively estimated at 36, 26,100 and 161,000 in HEK 293T; and 32, 19,000 and 124,000 in PGP1 iPSCs (Fig 1B). The TE’s copy number in HEK 293T is higher than that in PGP1 since the former cells are largely triploid. We used a complementary bioinformatic approach as a second estimate of TE abundance by aligning our designed gRNAs to the human reference genome (Fig 1C and *fig. S1*). An example of gRNA HL1gR4 targeting LINE-1 ORF2 is shown in Fig 1C. The total number of matches for HL1gR4 allowing 2 bp mismatches is 12,657, about half of our qPCR estimate, with the vast majority having an intact PAM (Fig 1D). Since the reference sequence likely undercounts TEs because of the well-known problems of assembling, aligning, and mapping these sequences(*28*), we base our editing numbers based on the qPCR copy number estimate.

**Figure 1.**
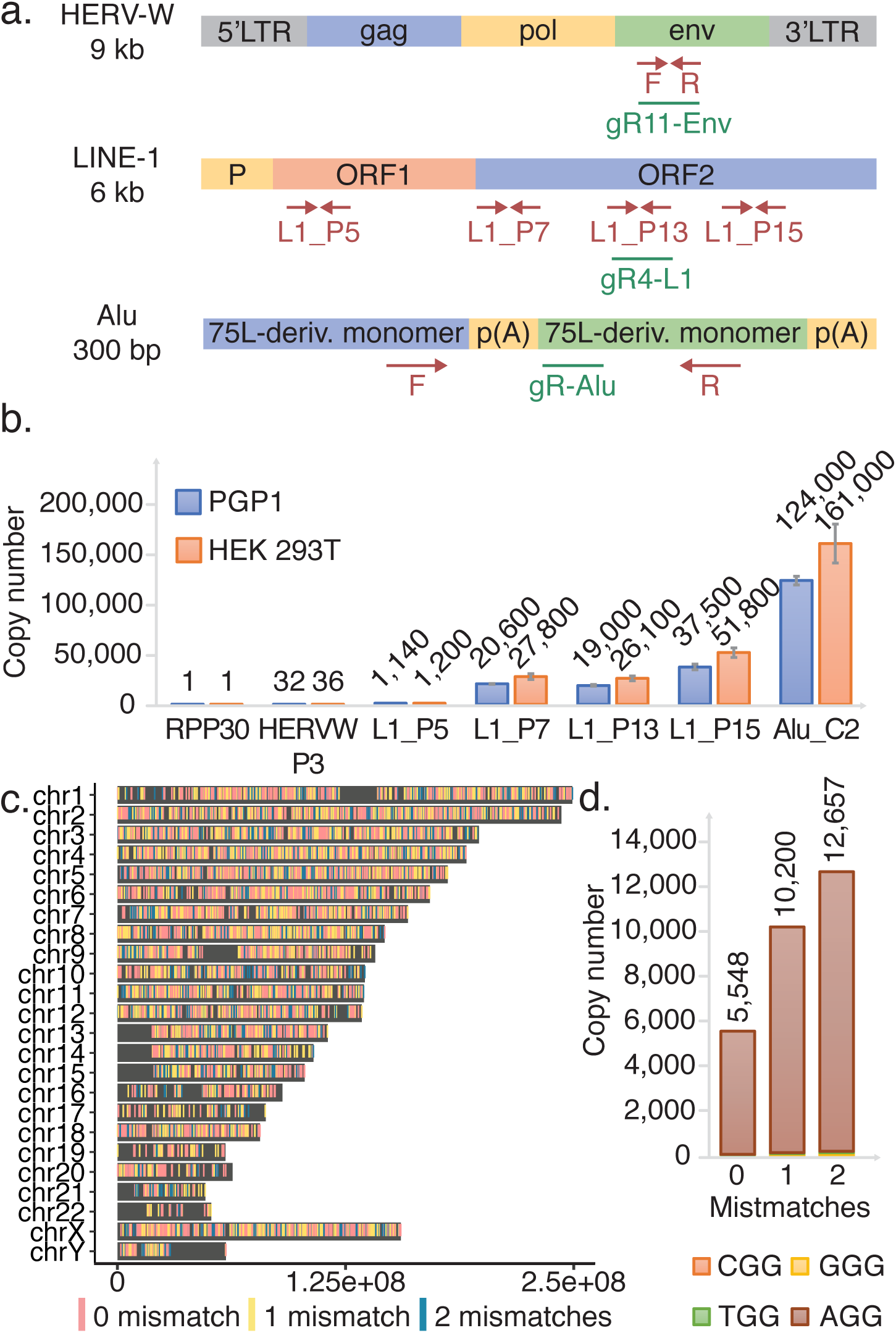
Utilizing high copy repetitive elements for the development of an extremely safe DNA editor. **(A)** A summary of HERV, LINE-1, and Alu. Representation of TEs with qPCR primer sites shown in red and gRNAs shown in green. **(B)** qPCR estimation of LINE-1 copy number across the element compared to single copy number controls in PGP1 and HEK 293T. Errors bars display standard deviation, n=3. (**C**) Genome wide distribution of HL1gR4. **(D)** HL1gR4 copy number and PAM distribution.

## High copy-number CRISPR/Cas9 editing induces cellular toxicity and inhibits survival of edited cells

We transfected HEK 293T cells with plasmids expressing pCas9_GFP and LINE-1 targeting gRNAs to disrupt the two key enzymatic domains of ORF-2: endonuclease (EN) and reverse transcriptase (RT) (Fig. 2A and *table S4*). Three days after transfection, we observed indel frequencies at the LINE-1 expected targets ranging from 1.3% to 8.7%, corresponding to an average of respectively 339 and 2271 edits per haploid genome in the population (Fig. 2B). In accord with previous reports that this degree of genetic alteration is toxic, we confirmed ~7-fold increases in cell death and apoptosis through Propidium Iodide and Annexin V staining (*fig. S3*). A follow-up time-course experiment provided evidence that cells that undergo editing at hundreds of loci do not survive. Here we transfected pairs of LINE-1 gRNAs targeting the EN, RT or both (ENRT) domains. Using pairs of gRNAs causes large deletions (~170-800 bp) that can be detected through gel visualization (*fig. S2A*). While samples from day two through five show clear editing with the expected deletion band sizes (Fig. 2C), they were no longer detectable at days 9 and 14 indicating that mutated cells either died out as suggested by our previous cell death assay or were overgrown by wild type cells. Deep sequencing of expected dual gRNA deletion bands confirmed the LINE-1 gRNA breakpoints (*fig. S2B*). While there were no visible bands at day 9 and 14, we repeated this experiment and attempted to isolate clones. After early indications of editing no clones had detectable mutations at day 12 and beyond (data not shown) suggesting that any significant level of indel activity at LINE-1 is toxic or limits growth and clonal isolation. Single cell analysis confirmed the bimodal editing frequency(*16*) with a mean deletion frequency of 47.1% (*fig. S4*).

**Figure 2.**
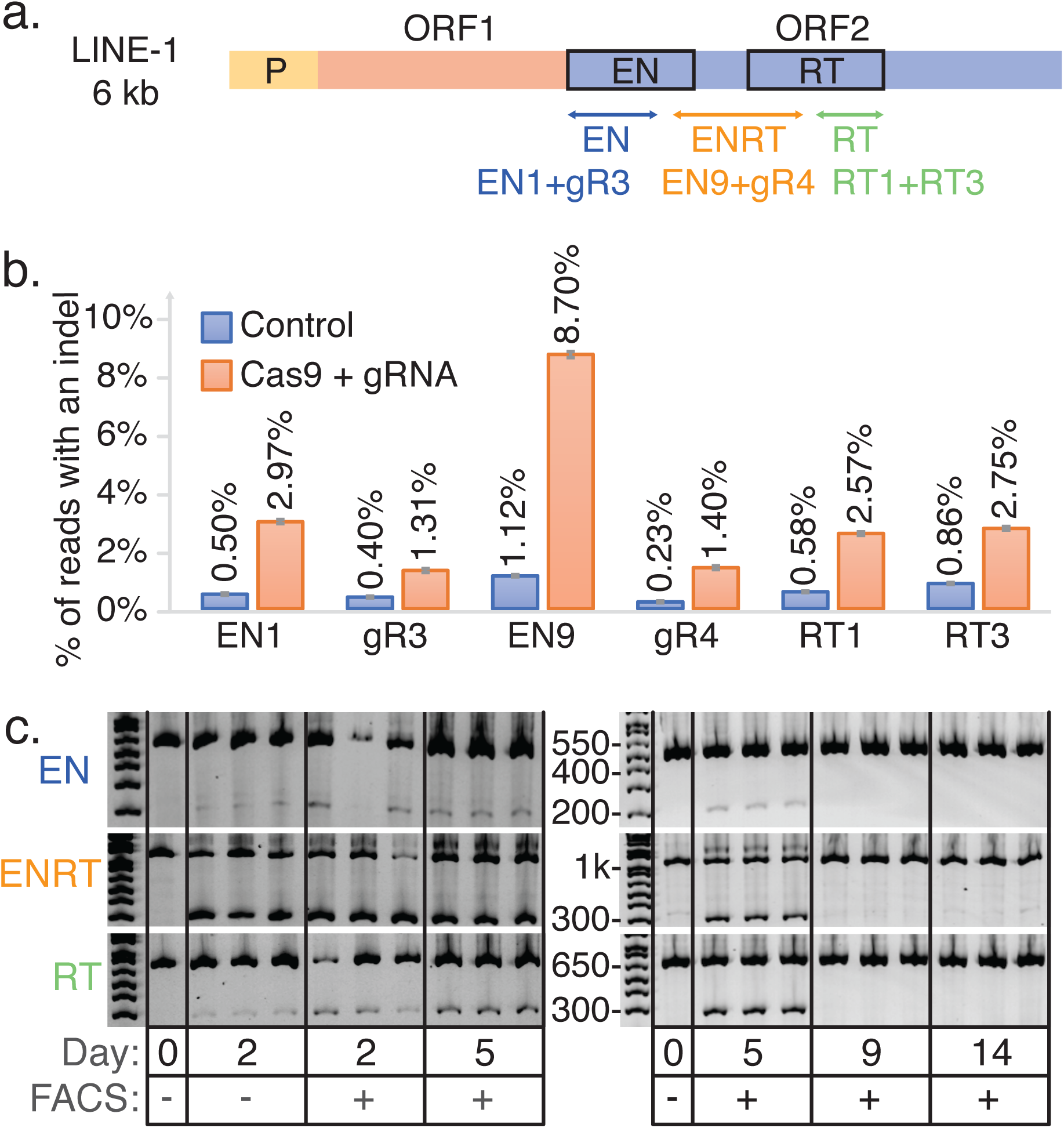
CRISPR-Cas9 based genome editing at high copy number repetitive elements is detectable but ultimately lethal. **(A)** Schematic of LINE-1 including the two protein coding genes ORF-1 and ORF-2. Three dual gRNA deletions were designed to disrupt the EN and RT domains of ORF-2. **(B)** LINE-1 gRNAs transfected with Cas9. Displayed are single transfections with 95% confidence intervals for a proportion as the error bars. **(C)** Gel image visualizing dual gRNA deletion bands compared to wild type control bands.

## nCBE and nABE enable isolation of stable cell lines with hundreds of edits

With the thought that nBEs could help improve the viability of LINE-1 edited cells, we designed and tested LINE-1 targeting gRNAs (HL1gR1-6 [*table S1*]) that generate a STOP codon early in ORF-2 using C→T deamination. When we transfected HEK 293T cells with nCBE3 and each of these gRNAs, we observed levels of deamination at each target locus that, although small (~0.05% – 0.67%) exceeded levels in mock transfected control cells (*fig. S5*). These same CBE gRNAs could also be used with ABEs as they contain at least one adenine within their deamination window. Above control levels of base editing were observed in genomic DNA in 4/5 gRNAs for both nCBE (*fig. S5B*) and nABE (*fig. S5C*). While nABE with HL1gR6 exhibited the highest editing efficiency (4.94% or ~1290 loci) three days after transfection, we used HL1gR4 going forward because it had the highest signal-to-background ratio among the more efficient gRNAs. The HL1gR4 target window also contained three efficiently-coedited C’s, thus offering a clear signal of directed mutation. In another experiment, an Alu targeting gRNA showed increased cell survival when using nCBE3 compared to Cas9 (*fig. S6*), suggesting that nBEs may reduce toxicity when targeting high copy number sequences.

293Ts were transfected with HL1gR4 and either nCBE3 or nCBE4-gam with control samples receiving a non-targeting gRNA. Two days post-transfection, single cells displayed up to 53.9% C→T deamination, or an estimated 14,000 loci (Fig. 3A), in the highest edited single cell. nCBE3 had a significantly higher mean deamination frequency than nCBE4-gam at this early timepoint but could not form any stable clonal population to the day 30 timepoint, suggesting that nCBE4-gam increased overall cell viability more than nCBE3 when targeting high copy repeats. Four surviving cell lines were isolated with deamination frequencies up to ~1.37 % of LINE-1 or an estimated ~356 sites (Fig 3B). Data presented in Fig. 3C shows both the purity of the desired deamination products and the editing window. Clone K was the highest edited stable clonal population and its targeted C→T mutation frequency from day 11 to 30 was confirmed.

**Figure 3.**
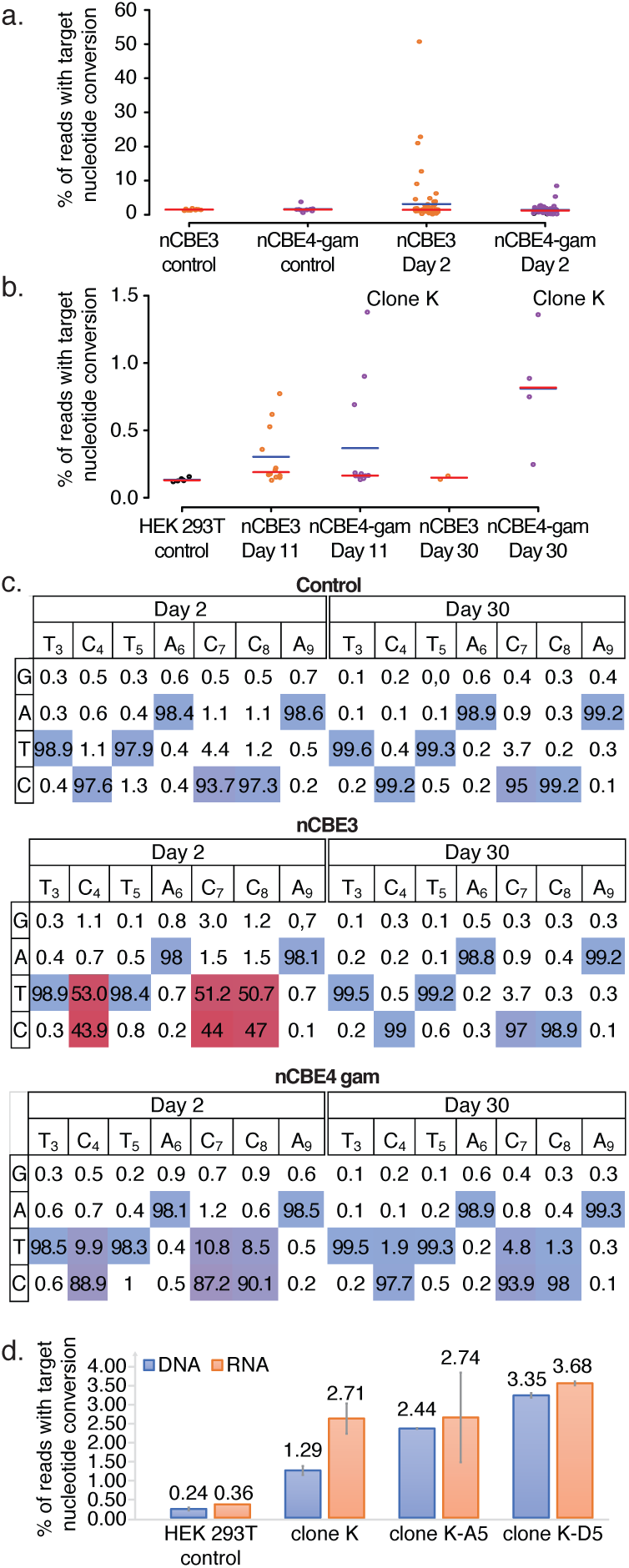
nBEs targeting LINE-1 enable survival of stable cell lines with hundreds of edits. **(A)** Base editing in HEK 293Ts two days after transfection comparing nCBE3 vs nCBE4-gam. FACS single cells are plotted as individual points representing targeted base editing nucleotide deamination. Red line indicates the median and the blue line the mean. **(B)** Single cell live culture growth and stable cell line generation at day 11 and 30. **(C)** Base editing activity across the CBE target window of ~3-9. Comparing day two and 30 for analysis of initial editing activity in most highly edited clones. **(D)** LINE-1 deamination analyzed from either RNA or genomic DNA. SEM’s are displayed as error bars, n=2.

By subjecting the top edited single cell isolate Clone K to another round of nCBE4-gam editing (*fig. S7A*) we detected cells with up to 36.26 % C→ T nucleotide conversion on day 2, and four living clones were isolated with mutation frequencies ranging from 2.43% to 5.04% – corresponding to about 643 to 1315 edits (*fig. S7B*). While the clone with the highest number of deaminated sites did not grow after a freezing and thawing cycle, the three other cell lines were stable in culture for a period longer than 30 days, and were termed “Clone K-A5”, “Clone K-A2” and “Clone K-D5”, with respectively 643, 749, and 781 edits, respectively. This observation of the highest edited clone dying off after initial detection was observed for all types of editors. We confirmed nBE activity at the lower copy number target HERV-W with up to 9.6% average nucleotide conversion at the population level (*fig. S8*). Due to the difficulty amplifying and analyzing the Alu target, likely because of high subfamily polymorphism and short repeat sequence (290bp) we proceeded exclusively with LINE-1 targeting gRNAs for the rest of the study.

To confirm that LINE-1 editing at the genome level was reflected on the corresponding transcripts we performed RNA-seq on Clone K, Clone K-D5, and Clone K-A5 and analyzed the percentage of C→T conversion resulting in a stop codon in ORF2 in the RNA reads (Fig. 3D). Theoretically, since most of the active LINE-1 subsets should generate transcripts, the presence of the expected stop codon at the messenger RNA level may indicate their inactivation. The results showed that a higher number of edits in the clones was correlated with a higher number of stop codons at the RNA level, suggesting that transcriptionally active LINE-1 subfamilies were impacted by the multiplexed editing.

## Nick-less dBEs enable the isolation of stable cell lines harboring up to 13,200 edits

Suspecting that generating single-stranded nicks genome-wide could lead to cytotoxicity, we decided to inactivate the remaining HNH nuclease domain of nCas9 by an H840A mutation in the nCas9 backbone and created a set of dCas9-BEs including dCas9-CBE4-gam (dCBE4-gam), dCas9-CBE4 (dCBE4), and dCas9-ABE (dABE). Nick-less dCas9-BEs were tested on single-locus targets to confirm their deamination activity and compare them to their nBE equivalents and the existing dCas9-CBE2 (dCBE2). dCBE4 and dCBE4-gam showed a 2.38- and 2.29-fold improvement in editing efficiency over CBE2 in 293Ts at day five respectively (*fig. S9A*). Compared to their nicking counterparts this was a 34.7% or 53.2% reduction in efficiency, but indel activity was reduced to background levels (*fig. S9A*). dABE retained 40.2% of nABE’s efficiency at a single locus target while reducing indel levels to background (*fig. S9B*).

We then transfected 293T cells with HL1gR4 and either nCBE4-gam, dCBE4-gam, nABE, or dABE and individually sorted and analyzed the cells for target nucleotide conversion after 2 days. Single edited cells resulted in editing efficiencies of up to 54.9% with nCBE4-gam, or 14,300 loci, while we observed significant reductions to mean target nucleotide mutation frequency with dCBE and dABE when compared to their nBE equivalents (Fig. 4A). In parallel, single cells were grown to determine whether viable highly edited clones could be isolated. The editing efficiency trend reversed in live cells: dBEs showed a significantly increased deamination frequency over nBEs (Fig. 4B). Remarkably, dABE produced the highest edited viable clone with 50.61% targeted nucleotide conversion or an estimated 13,200 loci. We estimate that, in our highest dCBE4-gam edited clone, we have inactivated 6292 of 26000 loci or 24.2% LINE-1 sequences. Base editors that retain nicking activity only generated a few rare cells with an editing frequency consistent with our prior experiments in Fig. 3B. Results were replicated using another LINE-1 targeting gRNA and similar trends were observed (*fig. S10*).

**Figure 4.**
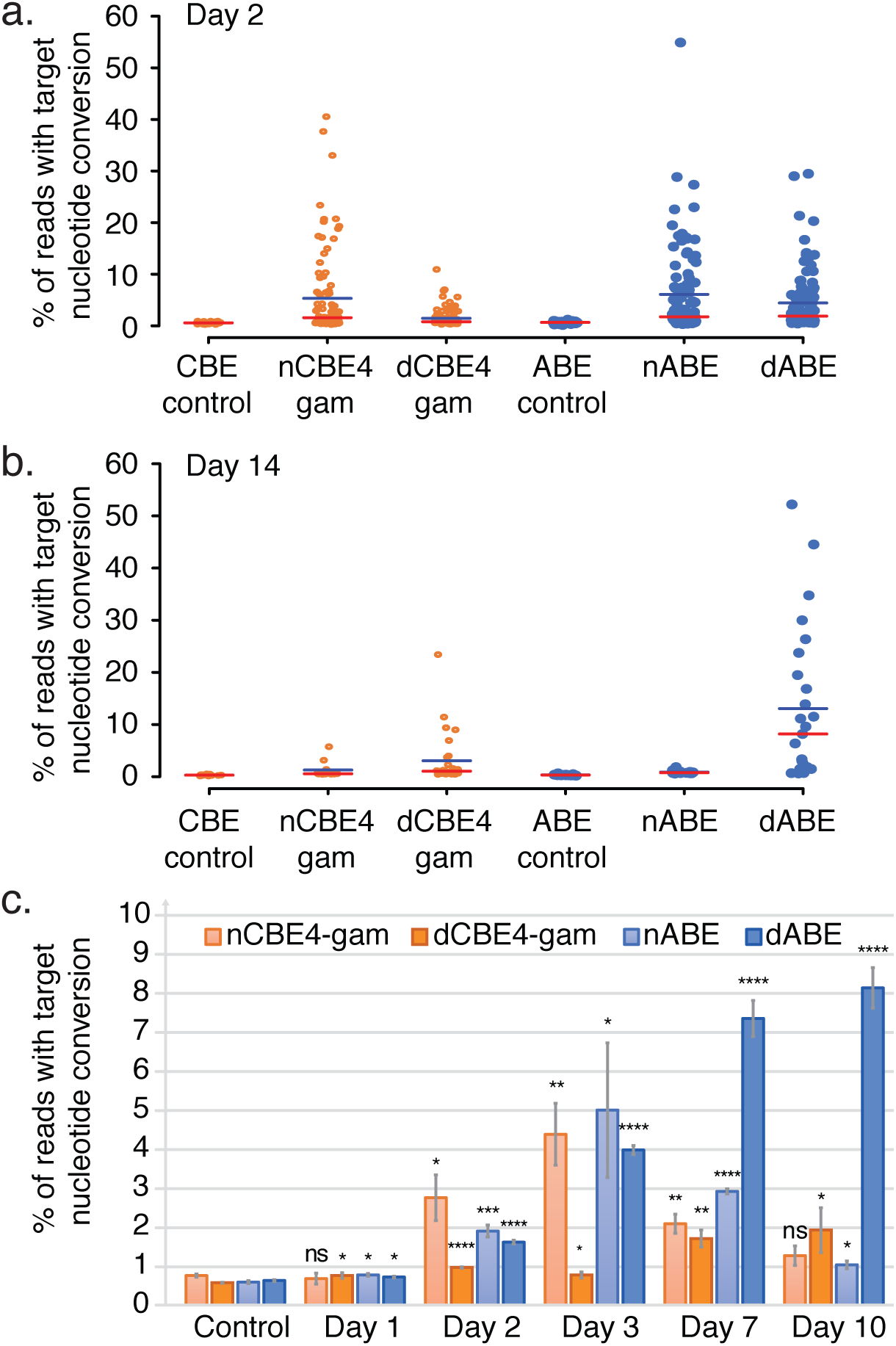
dBEs improve survival of highly edited cells with thousands of edits genome wide. **(A)** nBE compared to dBE in 293T single cells each represented as a single data point. Base editing is displayed as either target C->T or A->G conversion for CBE and ABE respectively. The red line indicates the median and the blue line the mean. **(B)** Live single cell analysis at day 14 of the same experiment. **(C)** Deamination frequency over time comparing dBE to nBE from day one to ten. Error bars represent SEM of n=3, ns (not significant), P ≥ 0.05; *P < 0.05; **P < 0.01; ***P < 0.001; ****P < 0.0001; by two-tailed Student’s t test compared to controls.

The nucleotide composition of all bases in the gRNA and PAM are displayed for the highest edited clone and parental 293T control for each BE condition used, showing very low non-specific nucleotide conversions for both nBEs and dBEs at LINE-1 (*fig. S11*). The mean single cell mutation frequency was reduced from 5.32% using nABE to 1.45% using dABE, indicating that disabling nicking resulted in a 3.67-fold decrease in editing efficiency at the day 2 timepoint (Fig. 4B). Fourteen days after transfection, dBEs gained a marked advantage as compared to nBEs in the total number of viable cells, and mutation frequency of single cells. There was a 14.8-fold increase in mean editing frequency among surviving live clones when using dABE compared to nABE (Fig. 4B), and a 2.38-fold increase was observed for dCBE4-gam compared to nCBE4-gam. High base editing purity was observed for both ABEs, while CBEs generated non-intended bases at the target position. dCBEs significantly reduced the generation of such non-intended bases at the target position, in particular dCBE4-gam (*fig. S9C and S9D*). No nonspecific nucleotide conversion was detected when targeting LINE-1 (*fig. S12*). During the first three days of editing the dBEs had lower editing frequency when compared to nBEs but from day seven, dABE gained a significant edge over nABE (Fig. 4C).

Chromosomal integrity analysis was performed for clones edited at LINE-1 with nABE, dABE, nCBE4-gam, and dCBE4-gam. The karyotype results are shown in *table S5* and show that the top edited clones are not significantly different from control groups in terms of the total number of aberrations (*fig. S13-S14*). Further analysis in a karyotypically normal and stable cell line is required to fully assess chromosomal stability after large-scale genome editing.

## dABE allows the isolation of hiPSCs harboring up to 2600 edits

We next attempted the large-scale genome editing of PGP1 hiPSCs. The survival cocktail and single cell isolation time line is shown in Fig. 5A. The same experiment was conducted with two slight variations of the electroporation protocol in terms of total cells transfected and the total amount of DNA used (CS and PK conditions). Single cells were sorted and analyzed for target nucleotide conversion frequency 18 hours post electroporation. The highest edited single cell had ~6.96% target A→G conversion or ~1320 sites (Fig. 5B). In parallel live single cells were isolated and stable cell lines formed at 11 days post transfection. Colonies were analyzed for targeted LINE-1 A→G nucleotide conversion with a 1.30% and 0.96% mean editing frequency for CS and PK conditions respectively 18 hours after transfection (Fig. 5C). At day 11, the median editing efficiency of the CS live clones was higher than that of PK live clones in contrast to the value observed at the earlier time point, suggesting that lower initial editing efficiency may increase the viability of stably edited cell lines. The highest edited clone had a nucleotide conversion frequency of 13.75% which corresponds to 2600 sites genome wide, exceeding by three orders of magnitude the number of simultaneous edits previously recorded in iPSCs(*29*). The increased background that occurs in single cell direct analysis Fig. 5B compared to isolation from an expanded colony Fig. 5C is likely due to the necessary over-amplification required to get enough genomic material from a single cell. Similar observations were made in previous experiments using 293T cells. All other previously tested DNA editors failed to produce any detectable edits at the LINE-1 locus in human iPSCs which are sensitive to even minor DNA damage(*30*) and rapidly deplete cells transfected with Cas9 and TE gRNAs over time (*fig. S15*).

**Figure 5.**
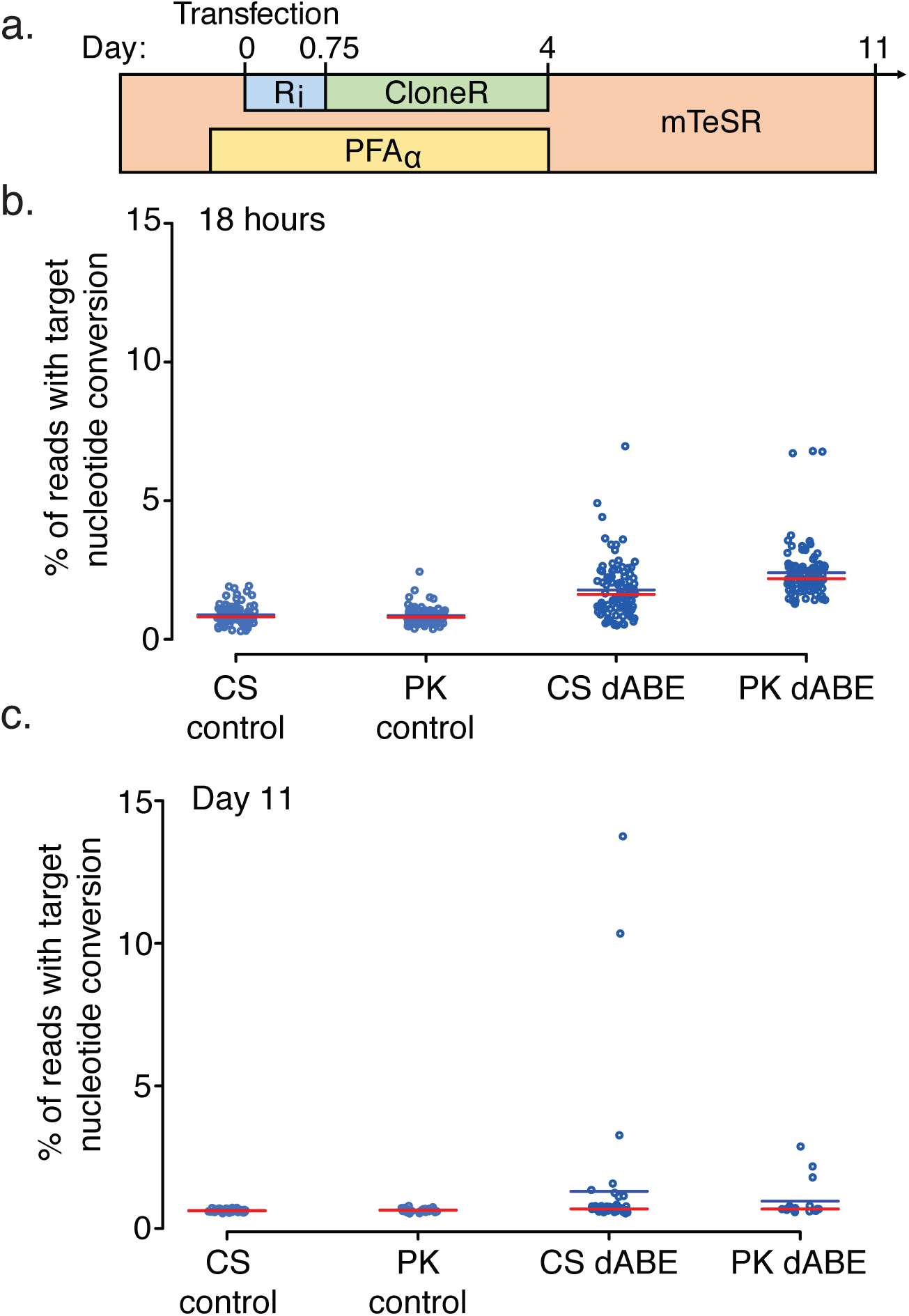
Survival cocktail and conditions for clonal derivation of iPSCs after large-scale genome engineering. **(A)** Human iPSC transfection timeline and survival cocktail conditions. **(B)** Eighteen-hour single cell direct NGS analysis of dABE targeting LINE-1. CS and PK indicate the two researchers who conducted the experiments. The red line indicates the median and the blue line the mean. **(C)** Live cell colony analysis of surviving iPSCs at day 11 post transfection.

## Discussion

CRISPR has recently brought about a radical transformation in the basic and applied biological research, leading to commercial applications and a multitude of clinical trials(*31*), and even to the controversial tests of human germline modifications(*32*–*36*). While the use of CRISPR and its myriad derivatives has greatly reduced the activation energy and technical skill required to perform genome editing several barriers limit fundamental and clinical applications: 1) The need for a custom gRNA, for each target, 2) difficult delivery, 3) inefficiencies once delivered, 4) off-target errors, 5) on-target errors, 6) the cytotoxicity of DNA damage when multiplexing beyond 62 loci(*16*), 7) the limitation of insertion to sizes below 7.4kb(*37*), 8) immune reactions to Cas, gRNA and vector. This study aims to develop tools that address weaknesses 5 and 6.

Improving the actual multiplexed eukaryotic genome editing capabilities by several orders of magnitude holds the potential of revolutionizing human healthcare. Combinatorial functional genomic assays would enable the study of complex genetic traits with applications in evolutionary biology, population genetics, and human disease pathology. Multiplex editing has also permitted the development of successfully engineered cell treatments, such as the chimeric antigen receptor (CAR) therapies, which require the simultaneous editing of three target genes. Future treatments may require many more modifications to augment cancer immunotherapies, slow down oncogenic growth, and reduce adverse effects, such as host-versus-graft disease. Furthermore, customizing host-versus-graft antigens in human- or nonhuman-donor tissues may require more modifications than have been made so far, for which the development of genome-wide editing technologies is needed. Special attention should be paid to the safety of the editing and its impact on the functional activity of the transplants, since donor tissues may persist in the patients for decades.

To complete genome-wide recoding and enable projects such as GP-write ultra-safe cells(*23*) or the de-extinction efforts to regain lost biodiversity, safe DNA editors must be developed to increase the number of genetic modifications possible by several orders of magnitude without triggering overwhelming DNA damage, as well as overcoming the delivery of multiple distinct gRNAs per cell, the latter of which we do not address is this study. C321.ΔA is a massively modified strain of *E. coli* MG1655 for which all instances of the Amber stop codon were replaced(*38*). To attempt such a feat in the human genome will require the modification of 4438 Amber codons(*24*). We have shown that gene editors that do not cause double- or single-strand DNA breaks can generate a number of edits sufficient to theoretically achieve this genome recoding and pave the way towards making pan-virus resistant human cells. This could have commercial application towards cell-based production of monoclonal antibodies, recombinant protein therapeutics, and synthetic meat production.

As our study demonstrates, genome-wide disruption of high copy number repetitive elements is now possible and opens new opportunities to study the “dark matter” of the genome. CBEs that allow the generation of STOP codons within an open reading frame will be a great tool to probe at the functions of transposable elements, potentially turning observed associations with physio-pathological phenotypes into causations. For instance, large-scale inactivation of HERV-W and LINE-1 elements could help investigate their respective roles in multiple sclerosis and neurological processes.

More in-depth studies will be necessary, however, to assess the impact of this massive editing on normal cell processes, since collateral damage may occur. We expect the thorough on-and off-target analysis at repetitive elements to remain a difficult task to accomplish due to their high level of polymorphism; therefore, strong biological controls as well as new experimental and bioinformatics pipelines will be needed to overcome such a challenge.

In our study, we observed that dABE increases the viability of highly edited clones as compared to dCBE. This difference may be explained by two factors. First, when using HL1gR4, CBE has three target nucleotides within its deamination window as compared to one for ABE, and as a consequence, CBE converts three times more nucleotide than ABE, potentially causing additional cytotoxicity. Second, when using CBE, the uracil N-glycosylase actively catalyzes the removal of the deaminated cytosine, generating several nicks genome-wide that promote DNA damage and potential cell death. The conversion of adenosine into inosine using ABE may not be detected as efficiently by the DNA repair machinery, therefore increasing the viability of large-scale editing. Random genome-wide off-target SNVs have been reported for CBEs that appear to be independent of gRNA binding sites(*39*). Thus, we anticipate the conditional modulation of DNA repair processes such as mismatch repair or base excision repair – that trigger downstream single- and double-strand breaks in the genome – to further improve the extent of dBEs’ performance.

Finally, since dBEs do not generate direct breaks in the genome, they decrease indel frequency to background and may not trigger DNA sensors such as p53, while retaining about 34% to 53% targeted nucleotide conversion frequencies as compared to their nBE counterparts. As a consequence, successful genetic modifications with dBEs may not enrich for pro-oncogenic cells that have disrupted DNA-damage guardians as has been reported for Cas9(*40*). Even at low levels of multiplexing, this feature may promote dBEs as an essential tool for therapeutic applications such as gene therapies.

In summary, this work optimized large-scale genome editing to enable cell viability after the simultaneous editing of thousands of loci per single cell. The ability to safely edit many loci genome wide may facilitate the true potential of personalized medicine as we further develop our understanding of gene interactions and epistasis. We envision these new safe DNA editors to be combined with further improvements in the multiplex delivery of gRNAs to usher in a new phase of synthetic biology where it is possible to imagine recoding whole mammalian genomes. When combined with further modulation of DNA repair and pro-survival factors there may be no limit to the number of bases that can be modified in a single genome, opening up new avenues that previously were thought not possible. We have overcome the toxicity limitation that prevented large-scale genome editing in human iPSCs and have expanded the editing boundary by three orders of magnitude. The continued development of multiplex delivery along with non-toxic, high-efficiency DNA editors without DSBs or SSBs is paramount to the success of genome-wide recoding efforts to probe the inner workings of life, ultimately leading to the radical redesign of nature and ourselves.

## Acknowledgments

We would like to thank John Aach, Reza Kalhor, Erkin Kuru, Javier Fernandez, Gabriel Filsinger, Timothy Wannier and Benedikt Markus for helpful discussions in the development of safe large-scale genome engineering; and Susan Byrne, Bobby Dhadwar, Alex Ng, and Alex Chavez for plasmids, cell lines, reagents and scientific support throughout. In addition, we would like to acknowledge Enzo Tramontano, Nicole Grandi, Angela Corona and Maria Paola Pisano for their help providing background and bioinformatics materials and support on HERV-W sequences. We would like to thank David Liu, Alexis Komor, Nicole Gaudelli and Johnny Hu for sharing their knowledge and reagents for base editing.

## Authors contribution

C.S., O.C., H.M., and G.M.C. initiated the study; C.S., O.C., K.S., P.K., and V.V. designed the experiments; C.S., O.C., K.S., P.K., V.V., C.W., and A.H. performed the experiments; C.S., O.C., K.S., P.K., V.V, R.F., M.G., and S.G. analyzed the data; C.S. and O.C. wrote the paper with contributing input from K.S., P.K., V.V., C.W., A.H., H.M., and G.M.C.

## Data and materials availability

Key plasmids developed during this study have been submitted to Addgene: pSB700_HL1 gR4 (# 124450), dABE (# 124447) and dCBE4-gam (# 124449). All NGS data used for the figures and supplementary figures have been made available at SRA BioProject Accession #PRJNA515875 and #PRJNA518077 for 293T and PGP1 respectively.

