## Supplementary materials for "Enabling large-scale genome editing by reducing DNA nicking"

**This PDF file includes:**

Materials and methods
Figures S1 to S16
Tables S1 to S5
Supplementary results
Data and materials availability
Supplementary references

#### MATERIALS METHODS

***Transposable element gRNA design*** - gRNAs targeting Alu were designed by downloading the consensus sequence from repeatmasker (http://www.repeatmasker.org/species/hg.html). LINE-1 gRNAs were designed based on the consensus of 146 "Human Full-Length, Intact LINE-1 Elements" available from the L1base 2(*1*). HL1gR 1-6 were designed to generate stop codons from C->T deamination mutations. EN, RT and ENRT pairs of gRNAs were designed to create moderate size deletions (200-800bp) easily distinguishable from their wild type full-length forms by gel visualization. HERV-W gRNAs were designed based on the consensus sequence of the 26 sequences identified by Grandi et al.(*2*) that can lead to the translation of putative proteins.

***qPCR evaluation of copy number across repetitive element targeting gRNAs*** - The qPCR reactions were performed using the KAPA SYBR FAST Universal 2X qPCR Master Mix (Catalog #KK4602) according to the manufacturer's instructions. The LightCycler 96 machine from Roche was used to perform the qPCRs and the results were extracted using the LightCycler 96 SW 1.1 software. The following thermocycling conditions were used: "preincubation" stage = 95°C for 180 sec; "2-step cycling" stage: annealing = 95°C for 3 sec and elongation = 60°C for 20 sec; "Melting" stage = keep standard. The following primers were used to perform the qPCRs.

| Primer Name | Sequence (5'-3') | Target |
|---|---|---|
| ZY-JAK2-F | AGCAAGTATGATGAGCAAGC | JAK2 |
| SB-JAK2-R | AAAACAGATGCTCTGAGAAAGGC | |
| P1(b)_REBE_F | TAGGAACAGCTCCGGTCTACA | LINE-1 promoter |
| P1_REBE-ilu_R | AATGCCTCGCCCTGCTTCGG | |
| P5_REBE-ilu_F | CCAATACAGAGAAGTGCTTAAAGG | LINE-1 ORF1 |
| P5_REBE-ilu_R | CTTGGAGGCTTTGCTCATTTCT | |
| P7_REBE-ilu_F | CCCATCAGTGTGCTGTATTCAGG | LINE-1 ORF2 |
| P7_REBE-ilu_R | GGCCTTCTTTGTCTCTTTTG | |
| P13_REBE-ilu_F | AACAGGCTCTGAAATTGTGGC | LINE-1 ORF2 |
| P13_REBE-ilu_R | GCTGGCCTCATAAAATGAGTTAG | |
| P15_REBE-ilu_F | GTTCTGGCCAGGGCAATCAG | LINE-1 ORF2 |
| P15_REBE-ilu_R | CCTGAGACTTTGCTGAAGTTGC | |
| P3_HERVWenv_F | AATACCACCCTCACTGGGCT | HERV-W env |
| P3_HERVWenv_R | CAGATTGGAAACAAGAGGTCC | |
| Alu_C2_F | CTGTAATCCCAGCACTTTGG | Alu |
| Alu_C2_R | CTCCGCCTCCCGGGTTCA | |

***Bioinformatic alignment and copy number analysis*** – Fasta sequences of hg38 reference genome were downloaded from Ensembl (ftp://ftp.ensembl.org/pub/release-95/fasta/homo_sapiens/dna/). Alignment analysis of the gRNA sequences to all chromosomes was performed using the R library Biostrings v2.40.2 and plotted using the R library ggplot2 3.3.0.

***SpCas9 and gRNA plasmids used for genome editing*** – The following Cas9 plasmids were used: pCas9_GFP (Addgene #44719), hCas9 (Addgene #41815). Base editing plasmids used: pCMV_BE3 (Addgene #73021), pCMV_BE4 (Addgene #100802), pCMV_BE4-gam (Addgene #100806), ABE 7.10 (Addgene #102909). The gRNAs used in this study were synthesized and cloned as previously described(*3*). Briefly, two 24mer oligos with sticky ends compatible for ligation were synthesized from IDT for cloning into the pSB700 plasmid (Addgene Plasmid #64046).

***SaCas9 and gRNA plasmids used for genome editing*** – Cas9 plasmid: pX600-AAV-CMV::NLS-SaCas9-NLS-3xHA-bGHpA (Addgene #61592). Base editing plasmid: SaBE4-gam (Addgene #100809). The gRNAs used in this study were synthesized and cloned as previously described(*4*). Briefly, two 24mer oligos with sticky ends compatible for ligation were synthesized from IDT for cloning into the BPK2660 plasmid (Addgene Plasmid #70709).

***Maintenance and transfection of HEK 293T cells*** - HEK293T cells were obtained from ATCC with verification of cell line identification and mycoplasma negative results. They were expanded using 10% fetal bovine serum (FBS) in high-glucose DMEM with glutamax passaging at a typical rate of 1:100 and maintained at 37°C with 5% $CO_2$. Transfection was conducted using Lipofectamine 2000 (Thermofisher Catalogue # 11668019) using the protocol recommended by the manufacturer with slight modifications outlined below. 24 hours before transfection ~$1.0 \times 10^5$ cells were seeded per well in a 12-well plate along with 1 mL of media. A total of 2 μg of DNA and 2 μL of Lipofectamine 2000 were used per well. For Cas9 plasmids, the DNA content per well was 1 μg of pCas9_GFP mixed with 1 μg of gRNA-expressing plasmid. For BE plasmids, 1.5 μg of BE was mixed with 0.5 μg of gRNA plasmid. In the dBE vs nBE comparison used to generate figure 4, Pifithrin-α (10 ng/μl) from Sigma-Aldrich P4359 (source # 063M4741V, Batch # 0000003019) was added to the media 30 minutes before transfection and maintained in the first day media change.

***FACS Single cell direct NGS preparation*** – To quantify early genetic editing in cells transfected with Cas9/BE and gRNA expression plasmids, single cells were sorted and prepared as follows. Two days post-transfection, single cells were FACS-sorted into 96-well PCR plates containing 10 μL of QuickExtract™ DNA Extraction Solution (Epicentre Cat. # QE09050) per well and genomic DNA (gDNA) was extracted using the manufacturer's protocol. Briefly, the sorted plates were sealed, vortexed and heated at 65°C for 6 minutes then 98°C for 2 minutes. The NGS library was prepared as described later below.

***Single cell clonal isolation and sequence verification*** – Single cells were FACS-sorted into flat bottom 96-well plates containing 100 μL of DMEM with 10% FBS and 1% Penicillin/Streptomycin per well. Sorted plates were incubated for ~14 days until well-characterized colonies were visible, with periodic media changes performed as necessary. To extract gDNA, the cells were first detached using 30 μL TrypLE™ Express (Thermofisher Cat. # 12604021) and neutralized with 30 μL growth media. Then, 4 μL of the resulting cell suspension was transferred to 10 μL of QE. Genomic DNA was extracted according to manufacturer's protocol, as described previously.

***Nested PCR Illumina MiSeq library preparation and sequencing*** - Library preparation was conducted as previously described(*5*). Briefly, genomic DNA was amplified using locus-specific primers attached to part of the Illumina adapter sequence. A second round of PCR included the index sequence and the full Illumina adapter. All PCRs were carried out using KAPA HiFi HotStart ReadyMix (KAPA Biosystems KK2602) according to the manufacturer's thermocycler conditions. Libraries were purified using gel extraction (Qiagen Cat. # 28706), quantified using Nanodrop and pooled together for deep sequencing on the MiSeq using 150 paired end (PE) reads.

***NGS indel analysis*** – Raw Illumina sequencing data was demultiplexed using bcl2fastq. All paired end reads were aligned to the reference genome using bowtie2(*6*) and the resulting alignment files were parsed for their cigar string to determine the position and size of all indels within each read using a custom perl script (*https://github.com/CRISPRengineer/mutation_indel*). All indels that were sequenced in both the forward and reverse reads were summed across all reads and reported for each sample along with the total number of reads. Indels within a 30 bp window from the 5' start of the gRNA proceeding through the PAM and extending an additional seven bp's (for a 20bp gRNA) were counted and summed for each sample.

***Dual gRNA deletion frequency NGS analysis*** – Reads were analyzed for dual gRNA large deletions by detecting sequences in between the gRNAs to indicate the full length unedited (at least not dual gRNA-edited) and sequences beyond the normal wild type amplicon that only appear when the deletion has occurred to identify deletion reads. The custom perl script used for analysis is available at *https://github.com/CRISPRengineer/dual_gRNA*.

***NGS base editing deamination analysis*** – All paired end reads were aligned to the reference genome using bowtie2, and the resulting alignment files were converted to bam, sorted, indexed, and variant called using samtools(*7*). All SNV data within a 30bp window from the 5' start of the gRNA proceeding through the PAM and extending an additional seven bp's (for a 20bp gRNA) are reported to analyze the editing window and purity of editing. The custom perl script used for analysis is available at https://github.com/CRISPRengineer/deamination_report.

***Site directed mutagenesis to remove remaining nick from base editors*** – We deactivated the remaining nuclease domain of Cas9 from nCBE4 (Addgene #100802), nCBE4-gam (Addgene #100806), and pCMV-ABE7.10 (Addgene #102919). Agilent QuikChange XL Site-Directed Mutagenesis Kit (catalogue # 200517) was used with the following primer sequences:

SpCas9-fwd – TTTATCTGATTACGACGTCGATGCCATTGTACCCCAATCCTTTTTG
SpCas9-rev – CAAAAAGGATTGGGGTACAATGGCATCGACGTCGTAATCAGATAAA

***Propidium Iodide and Annexin V staining and FACS analysis***
Cells were dissociated with TrypLE, diluted in an equal volume of PBS, and then centrifuged at ~300g for 5 minutes at room temperature. We resuspended samples into 500µl PBS and half of the cells were pelleted for later gDNA analysis. The remainder was centrifuged and resuspended into 100µl of Annexin V Binding Buffer (ref #V13246) diluted into ultrapure water at a 1:5 ratio. Subsequently, we added 5µl of Alexa 647 Annexin V dye (ref #A23204) and incubated samples in the dark for 15 minutes. We then added 100µl of Annexin V Binding Buffer and added 4µl of Propidium Iodide (ref #P3566) diluted into the Annexin V Binding Buffer at a 1:10 ratio. Samples were incubated in the dark for another 15 minutes. Cells were washed with 500µl of Annexin V Binding Buffer and centrifuged again to be finally resuspended into 400µl of Annexin V Binding Buffer. All samples were filtered using a cell strainer and were run on the LSR 11 using a 70-µm nozzle. Analysis was conducted using FlowJo software.

***Karyotype analysis of LINE-1 dBE-edited 293T single cell clones*** – Stable HEK 293T edited isolated cell lines (BE4-gam, dBE4-gam, ABE and dABE) were expanded and karyotypically compared with the control groups and the wild type HEK 293T. Actively growing cells were passaged 1-2 days prior to sending to BWH CytoGenomics Core Laboratory. The cells were received by the core at 60-80% confluency. Chromosomal count, variances and abnormalities were investigated.

***RNA-seq analysis of LINE-1 edited living cell lines*** – The RNA of 293T LINE-1 edited clones (1.37%-3.4% deamination by nCas9-CBE4-gam editing) was extracted by treatment with TRIzol (ThermoFisher Scientific, cat-# 15596018) followed by Direct-zol RNA Kit (Zymo Research, cat # R2072), according to the manufacturer's instructions. All samples were prepared from biological duplicates; the parental culture was divided into two cultures and passaged once before extraction. 500 ng RNA of each of the samples, as quantified by Qubit (QubitTM RNA HS Assay Kit, ThermoFisher Scientific, cat-# Q32852), was used to prepare the libraries using an NEBNext Directional RNA Library Prep Kit (New England Biolabs, cat-# E7765S) in conjunction with the Poly(A) mRNA Magnetic Isolation Module (New England Biolabs, cat-# E7490), and following the manufacturer's instructions.

Deamination frequency in the RNA was analyzed using the standard deamination analysis pipeline used for genomic DNA. Read counts were generated by mapping reads to a human reference genome (GRCh38.p12, using the PRI version from www.genecodegenes.org) using STAR. Differential gene expression analysis was performed in EdgeR version 3.24.3: Lowly expressed genes with less than 2 counts per million in 2 or more samples were filtered out, the libraries were normalized using TMM normalization and differentially expressed genes were identified by using the exact test on the tagwise dispersion to compare the expression of each of the clones to the control sample. The Benjamini-Hochberg method was used to adjust p-values for multiple testing.

Multidimensional Scaling distances were generated by using the plotMDS function of EdgeR on the filtered and normalized libraries and plotted using ggplot.

***Maintenance and expansion of human iPSCs*** - Human iPSCs were cultured with mTeSR medium on tissue culture plates coated with Matrigel (BD Biosciences). For routine passaging, iPSCs were digested with TrypLE (Thermofisher # 12604013) for 5 minutes and washed with an equal volume PBS by centrifugation at 300g for 5 minutes. Digested iPSC pellets were physically broken down to form a single cell suspension and then plated onto Matrigel-coated plates at a density of $3x10^4$ per $cm^2$ with mTeSR$^{TM}$ medium supplemented with 10µM Y-27632 ROCK inhibitor ($R_i$) (Millipore, 688001) for the first 24 hours.

***Nucleofection in PGP-1 iPSCs*** – Thirty minutes prior to transfection media was changed to mTeSR$^{TM}$ supplemented with Pifithrin-α (10 ng/µl) from Sigma-Aldrich P4359 (source # 063M4741V, Batch # 0000003019); a notable spiky edge colony morphology was observed similar to when $R_i$ is added. Human iPSCs were digested with TrypLE for 5 minutes and the single cells were washed once with PBS. (CS: 4 x $10^6$, PK: 1 x $10^6$) iPSCs were then re-suspended in 100 µl of P3 Primary Cell Solution (Lonza) supplemented with (CS: 13.5 µg, PK: 6.75 µg) of dABE plasmid, (CS: 4.5 µg, PK: 2.25 µg) of gRNA plasmid, and (CS: 2 µg, PK: 1 µg) of pMax. The combined cells and DNA were then nucleofected in 4D-Nucleofector (Lonza) using the hES H9 program (CB150). The nucleofected iPSCs were then plated onto a single well of a 6-well Matrigel-coated plate in mTeSR medium supplemented with 10 μM $R_i$ and Pifithrin-α (10 ng/μl).

***Clonal isolation of PGP-1 iPSCs*** – 96-well plates were coated with Matrigel (BD Biosciences) at a concentration of 50 μl/well. A cloning medium solution of 10% CloneR™ (StemCell Technologies #05888) and Pifithrin- α (10 ng/μl) in mTeSR™ was prepared and added to the coated wells. Cells were digested using TrypLE, which was neutralized by an equal amount of cloning medium. The cell solution was then centrifuged at 300 x g for 5 minutes, the supernatant was aspirated, and the cell pellet was resuspended in the cloning medium. The cells were then passed through a 40-μm cell strainer and were FACS-sorted into 1) individual wells containing warm cloning medium at a density of 1 cell/well and 2) 2 x 96-well PCR plates for direct NGS analysis. To prevent disturbance, there was no media change during the first 48 hours, and the plates were not removed from the incubator during this period. A half-medium change was performed on days 3 and 4 with cloning medium. The growing colonies were monitored and a mTeSR™ medium change was done daily for the following days until extracting the DNA using QuickExtract™ and proceeding with library preparation and sequencing.

***Statistical Analysis*** –Statistical analysis was conducted using the student's t test. Differences were considered significant if p value was <0.05. * - 0.01 < p <0.05, ** - 0.001 < p <0.01, *** - p <0.001, **** - p <0.0001.

#### SUPPLEMENTARY FIGURES

**(A)**

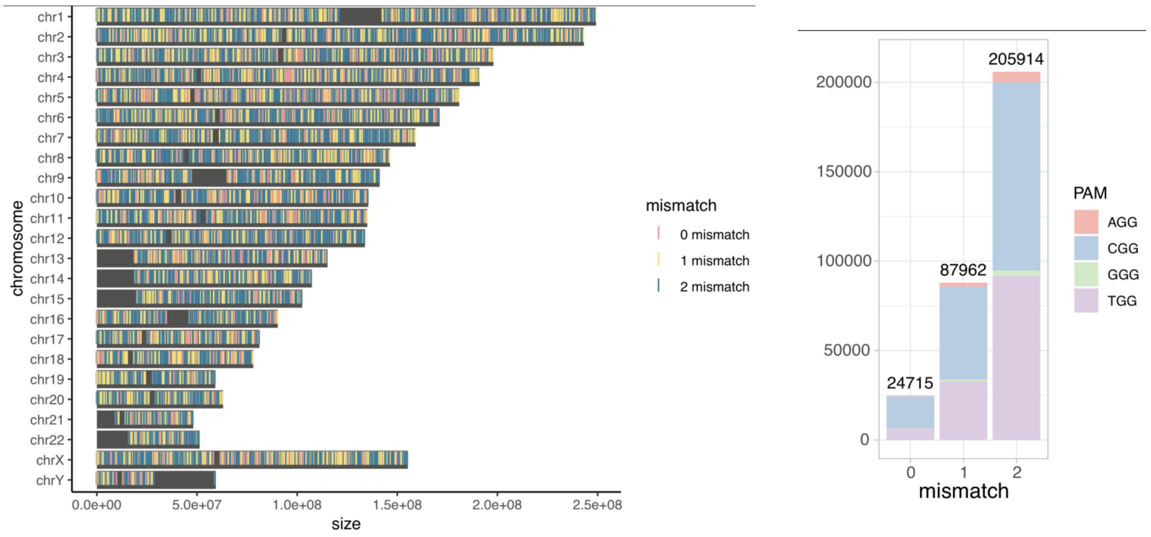


**(B)**

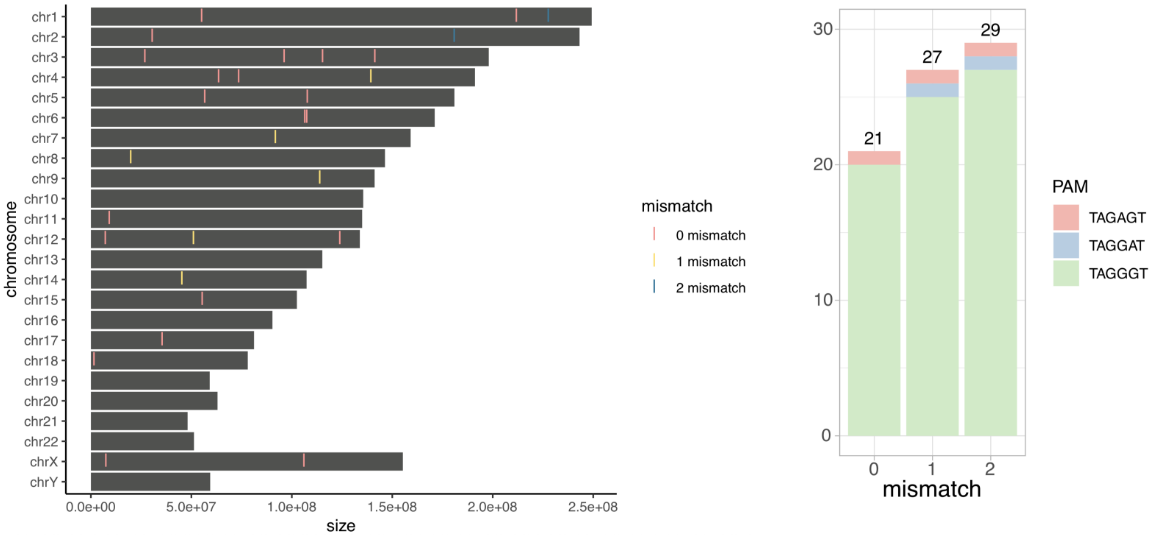


**Figure S1 | TE gRNA human reference alignment.** (**A**) (left) Genome wide distribution of gRNA Alu (right) Alu copy number and PAM distribution. (**B**) (left) Genome wide distribution of gRNA HERV env11 (right) HERV copy number and PAM distribution.

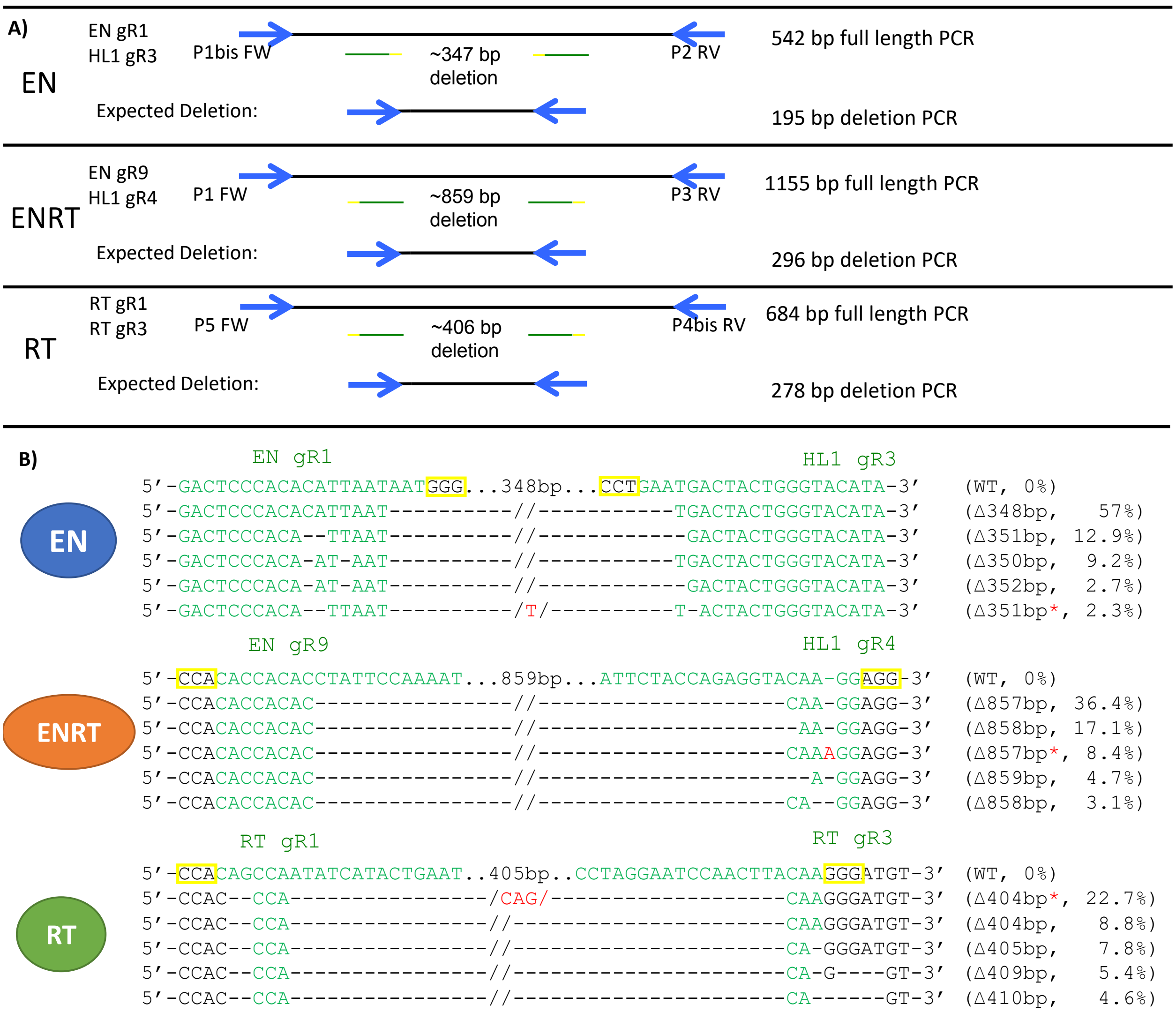


**Figure S2 | dual gRNA LINE-1 deletions.** (**A**) Primers used to amplify dual gRNA pairs targeting LINE-1 with full length and expected deletion product sizes shown. gRNAs are represented as green with yellow PAMs, primers are in blue. (**B**) Dual gRNA deletion frequency displaying the expected cut points near each gRNA. Green nucleotides are within the gRNA sequence, red are inserted nucleotides and "-" are deletions. The sizes of deletions and percentage among sequencing reads are displayed to the right.

**A)**

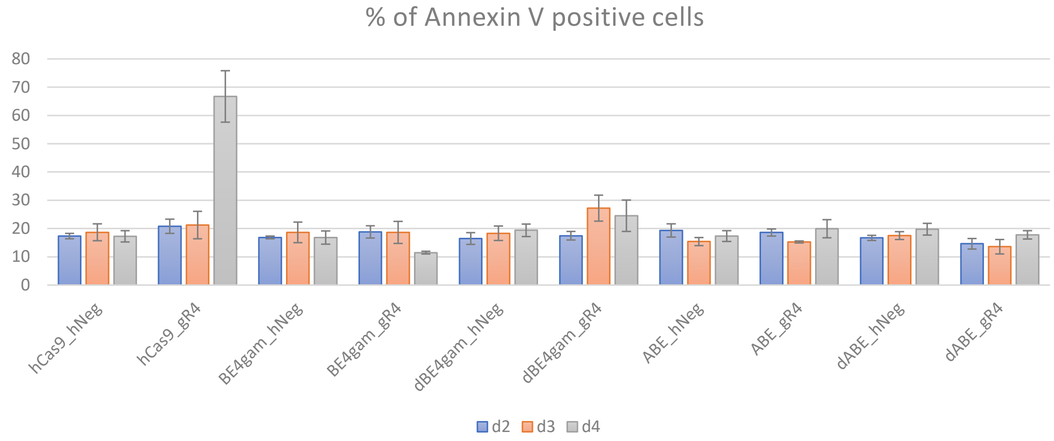


**B)**

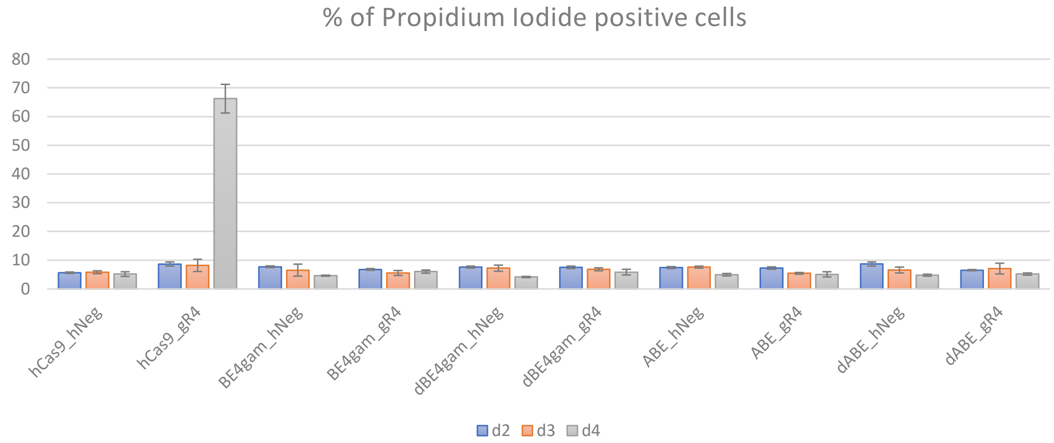


**C)**

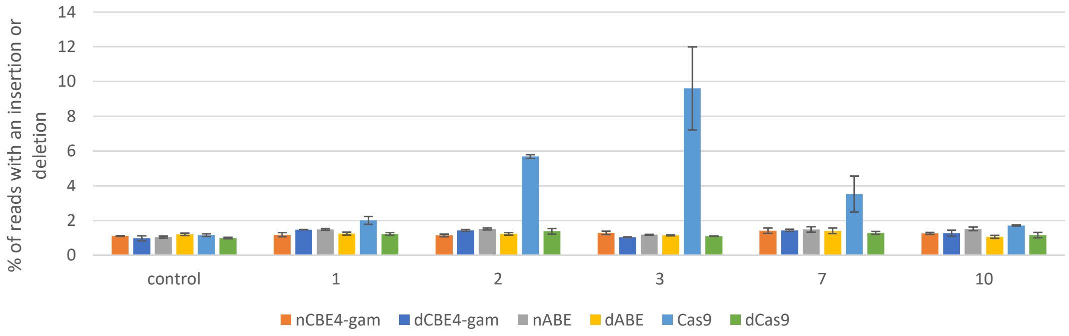


**Figure S3 | Annexin V and propidium iodide assays for cytotoxicity. (A)** Apoptosis cell death analysis using Annexin V targeting LINE-1. **(B)** Necrosis cell death analysis using propidium iodide. (**C**) Indel mutagenesis analysis with experimental day on the x-axis.

A)

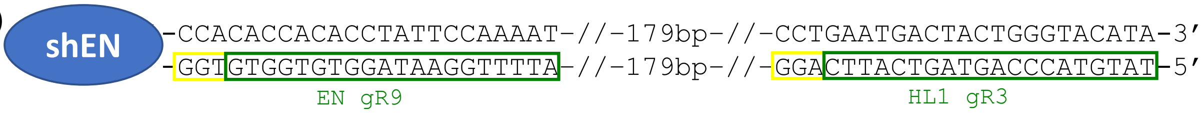


B)

| condition | total cells screened | # of edited cells | % of edited cells |
|---|---|---|---|
| All samples | 364 | 303 | 83.24 |

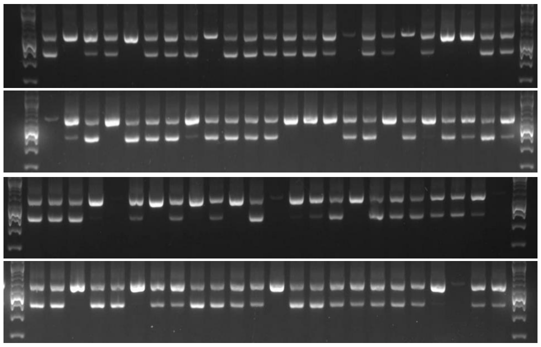

C)

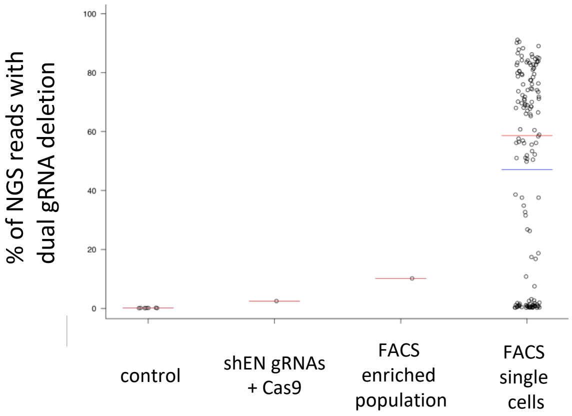


**Figure S4 |** Single cell analysis of dual gRNA deletions targeting LINE-1 **(A)** gRNA targets used for the shEN dual deletion. (**B**) Gel visualization of dual gRNA deletions bands in FACS single cells with a summary table. (**C**) Percentage of single cells with dual gRNA deletions.

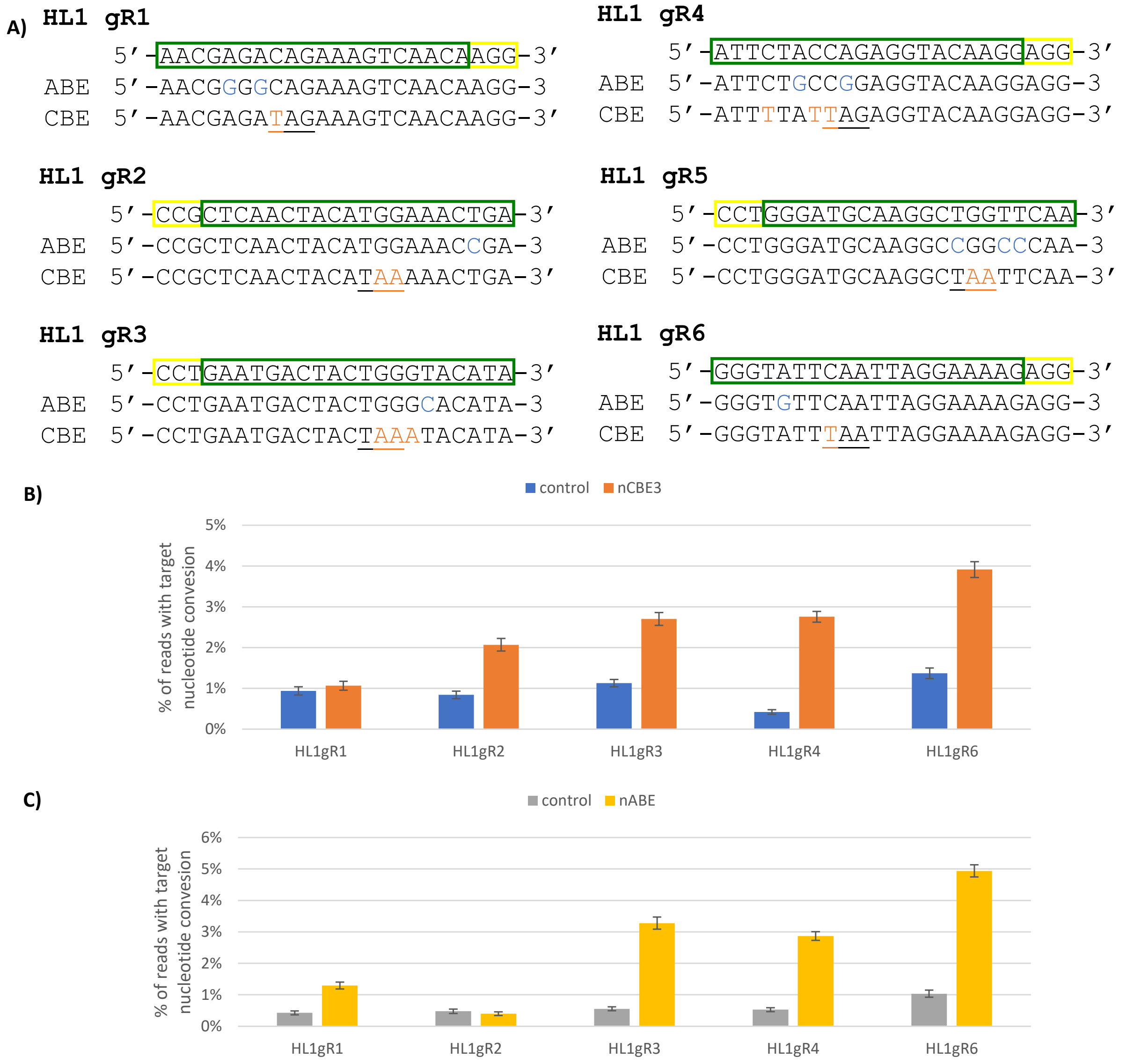


**Figure S5 | nBE targeting LINE-1 (A)** LINE-1 gRNA targets outlined in green boxes with PAMs in yellow. Expected ABE and CBE deamination products are displayed below with altered bases in blue and orange respectively **(B)** Targeted deamination frequency at C8 using nCBE3. **(C)** Targeted deamination frequency at A6 using nABE.

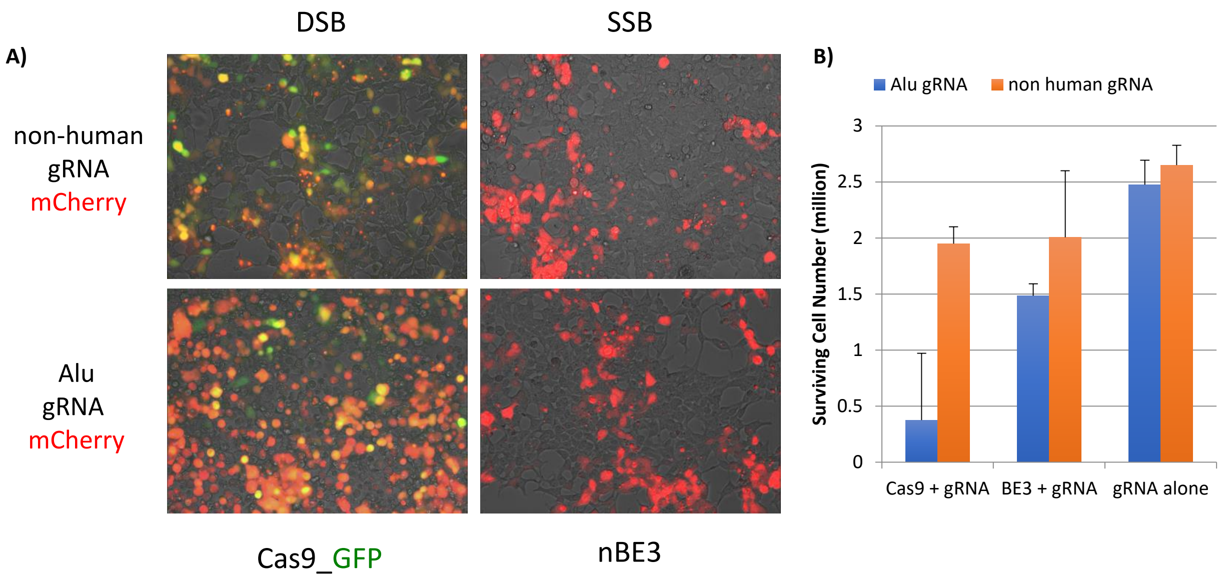


**Figure S6 | nCBE3 vs Cas9 targeting Alu in HEK 293Ts. (A)**Microscope images of rapid cell death in cells that express Cas9 along with a gRNA that targets a high copy number locus. HEK293T cells were transfected with a gRNA targeting the Alu consensus sequence along with either Cas9 that generates a DSB or nCBE3 which generates a single stranded break. Cells were imaged 72 hours after transfection. As a control a non-human targeting gRNA was used to determine the background survival after transfection under the same conditions. **(B)** Total cell count comparing the Alu gRNA in blue and the nonhuman gRNA in orange that was transfected with Cas9_GFP or no nuclease.

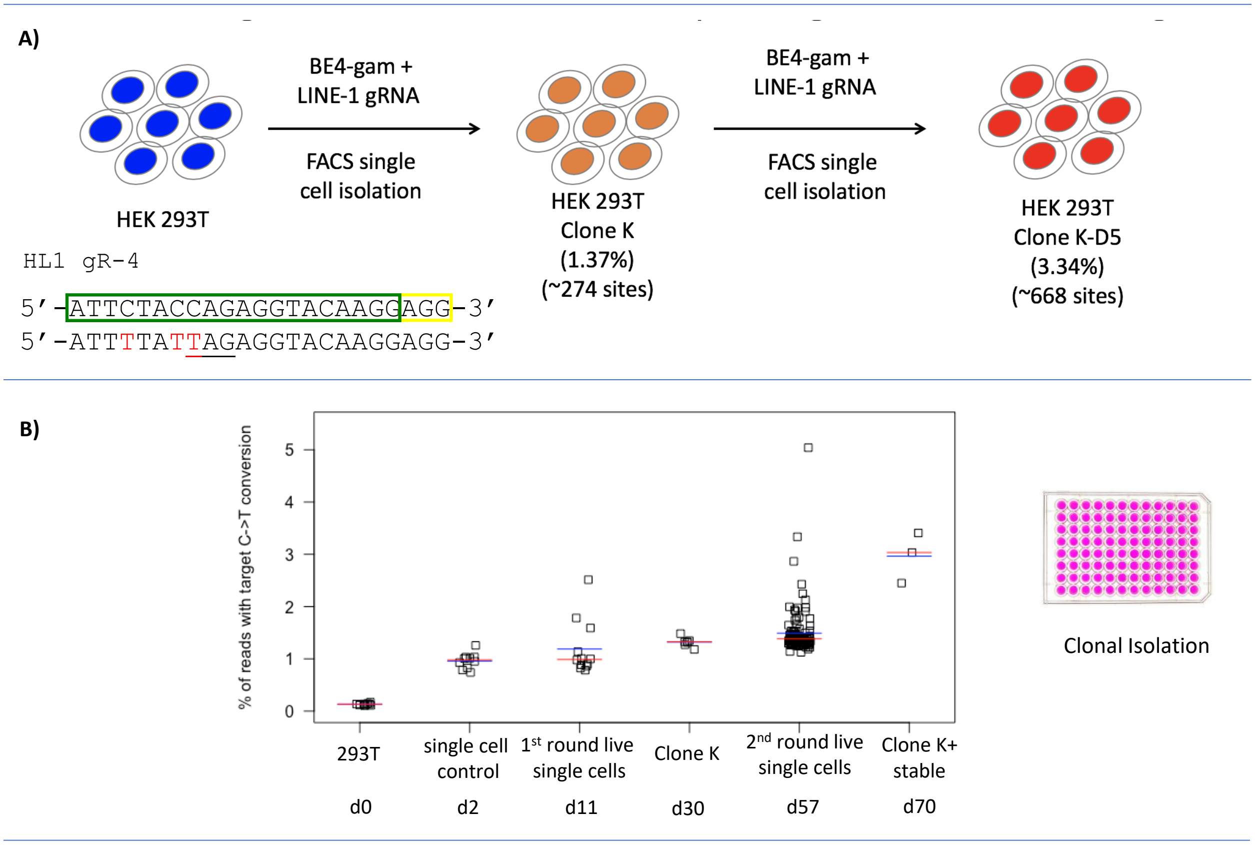


**Figure S7 | Utilizing high copy repetitive elements for the testing of an extremely safe DNA editor.** **(A)** Experimental design for two rounds of base editing at LINE-1. gRNA target is outlined in green with a yellow PAM. C->T deamination targets are colored in red. **(B)** Targeted deamination frequency at C8 using nCBE4-gam over two rounds of transfection and clonal isolation.

A)

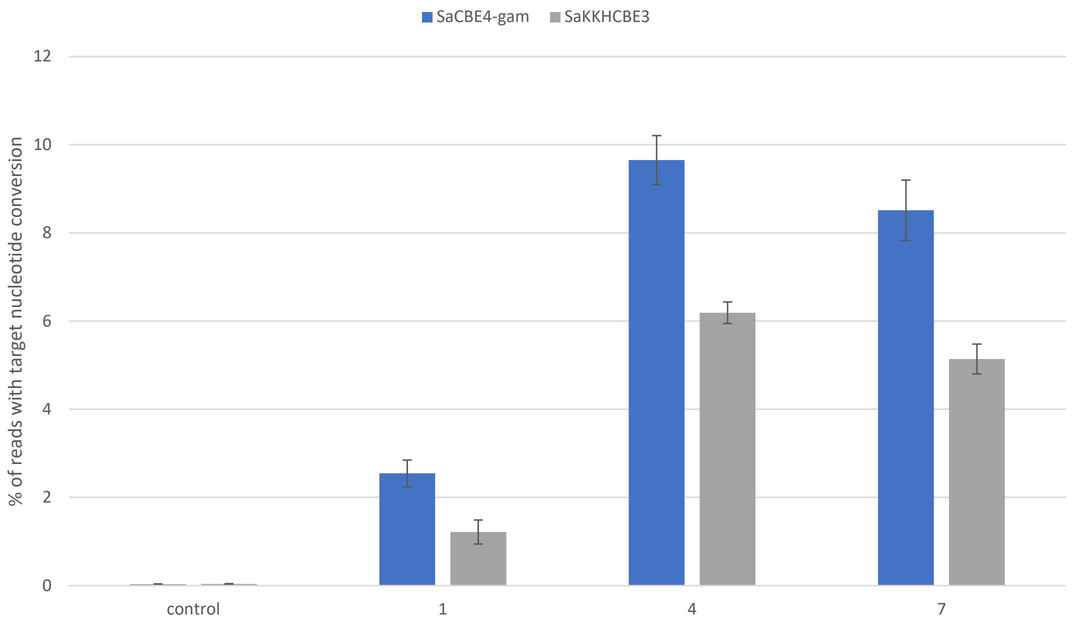


**Figure S8 | Base editing at HERV. (A)** Targeted deamination frequencies at C12 using a set of SaCas9-BEs over three time points (day 1, 4 and 7). Error bars represent SEM, n=3.

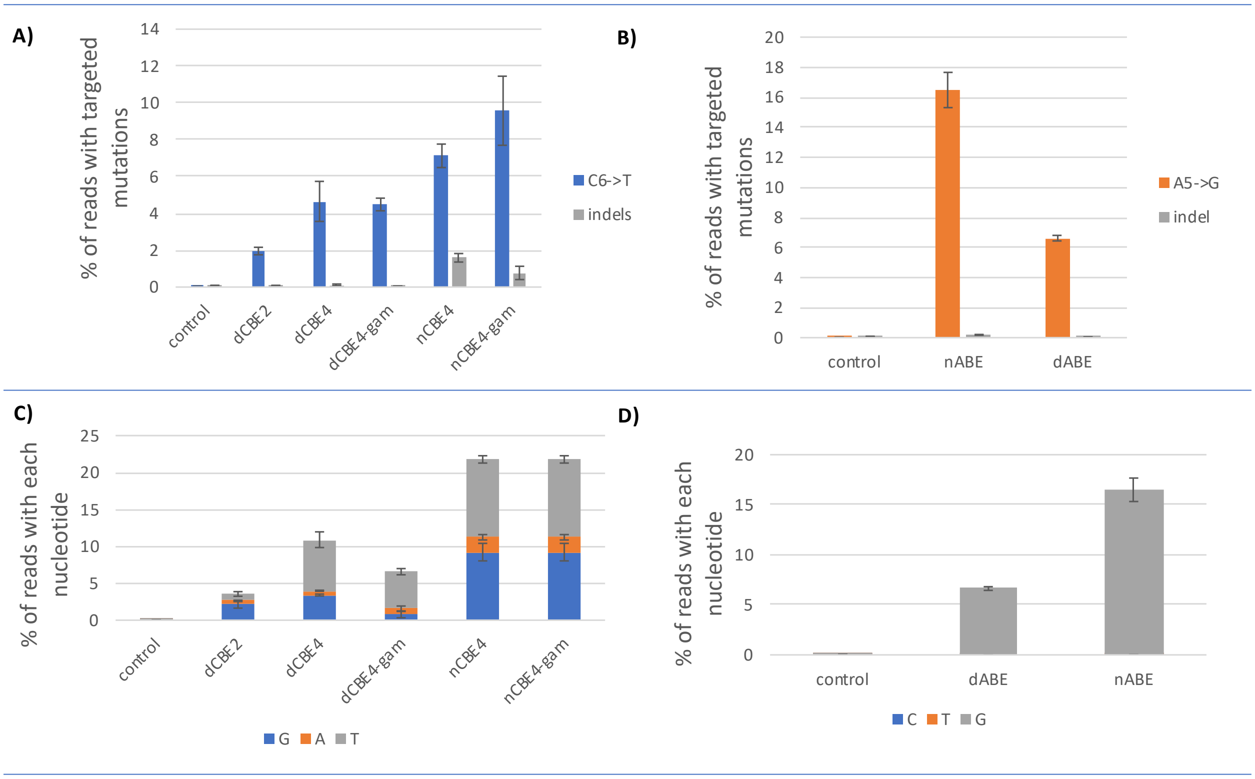


**Figure S9 | dBE vs nBE at a single locus target. (A)** Targeted deamination and indel frequencies at C6 using a set of CBEs and gRNA S1. Error bars represent SEM, n=3. **(B)** Targeted deamination and indel frequencies at A5 using ABEs.(**C**) Base editing purity analysis of C6. (**D**) Purity analysis for A5 using ABEs.

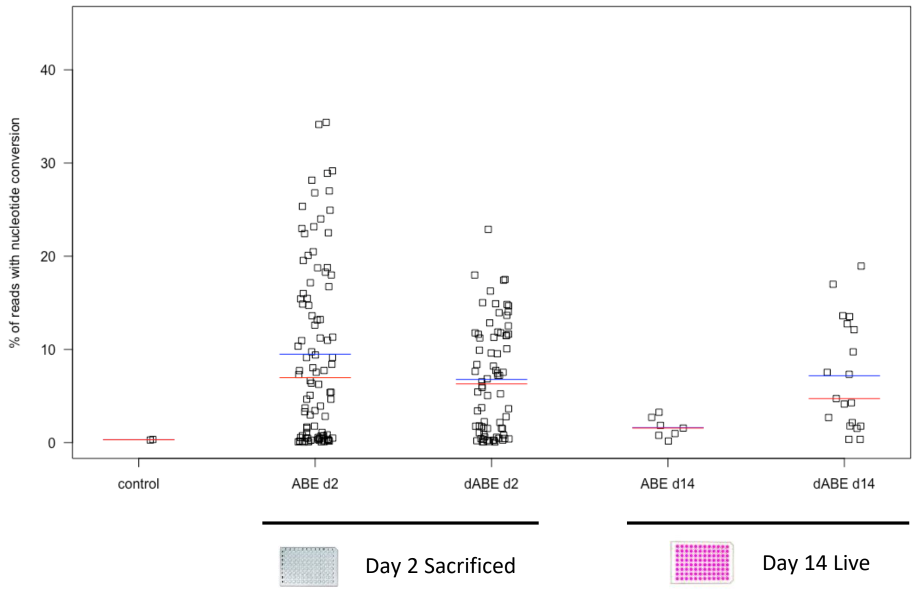


**Figure S10 | dABE targeting LINE-1 single cell analysis. (A)** Base editing in HEK 293Ts after transfection comparing nABE vs dABE with HL1gR46 at days 2 and 14. FACS single cells are plotted as individual points representing targeted base editing nucleotide deamination. Red line indicates the median and the blue line the mean.

**A)**

control

| | | | | | | | | | | | | | | | | | | | | | PAM | |
|---|---|---|---|---|---|---|---|---|---|---|---|---|---|---|---|---|---|---|---|---|---|---|
| | $A_1$ | $T_2$ | $T_3$ | $C_4$ | $T_5$ | $A_6$ | $C_7$ | $C_8$ | $A_9$ | $G_{10}$ | $A_{11}$ | $G_{12}$ | $G_{13}$ | $T_{14}$ | $A_{15}$ | $C_{16}$ | $A_{17}$ | $A_{18}$ | $G_{19}$ | $G_{20}$ | A | G | G |
| G | 0.3 | 0.1 | 0.2 | 0.2 | 0.1 | 0.6 | 0.3 | 0.4 | 0.3 | 98.1 | 0.3 | 98.3 | 97.7 | 0.1 | 1.0 | 1.0 | 0.7 | 0.6 | 96.9 | 97.7 | 0.4 | 97.7 | 98.6 |
| A | 99.2 | 0.2 | 0.2 | 0.2 | 0.1 | 98.6 | 0.9 | 0.4 | 99.2 | 0.5 | 99.1 | 0.9 | 1.2 | 0.1 | 98.6 | 1.2 | 98.8 | 97.8 | 2.8 | 1.4 | 99.4 | 1.8 | 0.7 |
| T | 0.3 | 99.7 | 99.3 | 0.6 | 99.0 | 0.2 | 3.5 | 0.4 | 0.3 | 0.2 | 0.3 | 0.4 | 0.7 | 99.1 | 0.2 | 1.9 | 0.2 | 1.5 | 0.3 | 0.4 | 0.1 | 0.4 | 0.4 |
| C | 0.1 | 0.3 | 0.3 | 98.9 | 0.7 | 0.4 | 95.1 | 98.7 | 0.1 | 1.0 | 0.2 | 0.6 | 0.4 | 0.6 | 0.1 | 96.1 | 0.3 | 0.2 | 0.1 | 0.5 | 0.1 | 0.2 | 0.3 |

nCBE4-gam

| | | | | | | | | | | | | | | | | | | | | | PAM | |
|---|---|---|---|---|---|---|---|---|---|---|---|---|---|---|---|---|---|---|---|---|---|---|
| | $A_1$ | $T_2$ | $T_3$ | $C_4$ | $T_5$ | $A_6$ | $C_7$ | $C_8$ | $A_9$ | $G_{10}$ | $A_{11}$ | $G_{12}$ | $G_{13}$ | $T_{14}$ | $A_{15}$ | $C_{16}$ | $A_{17}$ | $A_{18}$ | $G_{19}$ | $G_{20}$ | A | G | G |
| G | 0.3 | 0.0 | 0.1 | 0.5 | 0.1 | 0.6 | 0.4 | 0.5 | 0.3 | 98.1 | 0.3 | 98.4 | 97.9 | 0.1 | 1.1 | 0.7 | 0.6 | 0.5 | 96.8 | 98.2 | 0.3 | 97.9 | 98.2 |
| A | 99.3 | 0.1 | 0.1 | 0.2 | 0.1 | 98.7 | 0.9 | 0.4 | 99.2 | 0.6 | 99.3 | 0.8 | 1.2 | 0.1 | 98.5 | 1.2 | 99.0 | 97.9 | 2.9 | 1.1 | 99.4 | 1.7 | 0.9 |
| T | 0.2 | 99.7 | 99.5 | 7.4 | 99.3 | 0.2 | 9.9 | 6.9 | 0.3 | 0.1 | 0.3 | 0.3 | 0.6 | 99.2 | 0.3 | 2.1 | 0.1 | 1.4 | 0.2 | 0.2 | 0.2 | 0.3 | 0.5 |
| C | 0.2 | 0.2 | 0.2 | 91.8 | 0.4 | 0.4 | 88.7 | 92.2 | 0.1 | 1.1 | 0.1 | 0.3 | 0.2 | 0.6 | 0.1 | 96.0 | 0.2 | 0.2 | 0.1 | 0.5 | 0.1 | 0.2 | 0.4 |

dCBE4-gam

| | | | | | | | | | | | | | | | | | | | | | PAM | |
|---|---|---|---|---|---|---|---|---|---|---|---|---|---|---|---|---|---|---|---|---|---|---|
| | $A_1$ | $T_2$ | $T_3$ | $C_4$ | $T_5$ | $A_6$ | $C_7$ | $C_8$ | $A_9$ | $G_{10}$ | $A_{11}$ | $G_{12}$ | $G_{13}$ | $T_{14}$ | $A_{15}$ | $C_{16}$ | $A_{17}$ | $A_{18}$ | $G_{19}$ | $G_{20}$ | A | G | G |
| G | 0.3 | 0.0 | 0.0 | 1.1 | 0.0 | 0.3 | 0.6 | 0.2 | 0.2 | 97.7 | 0.1 | 98.6 | 98.3 | 0.0 | 0.7 | 1.4 | 0.6 | 0.6 | 96.3 | 97.8 | 0.4 | 97.6 | 98.6 |
| A | 99.5 | 0.0 | 0.1 | 0.2 | 0.1 | 98.5 | 1.3 | 0.4 | 99.1 | 0.7 | 99.4 | 0.6 | 1.1 | 0.1 | 98.7 | 1.3 | 99.3 | 97.3 | 3.3 | 1.7 | 99.4 | 1.9 | 0.9 |
| T | 0.1 | 99.7 | 99.5 | 23.9 | 99.0 | 0.2 | 25.8 | 24.2 | 0.3 | 0.3 | 0.1 | 0.3 | 0.4 | 99.2 | 0.3 | 2.2 | 0.2 | 1.8 | 0.3 | 0.1 | 0.2 | 0.5 | 0.2 |
| C | 0.0 | 0.2 | 0.1 | 74.3 | 0.5 | 0.4 | 71.8 | 74.9 | 0.0 | 0.9 | 0.1 | 0.3 | 0.2 | 0.4 | 0.1 | 96.4 | 0.1 | 0.3 | 0.0 | 0.3 | 0.0 | 0.1 | 0.3 |

nABE

| | | | | | | | | | | | | | | | | | | | | | PAM | |
|---|---|---|---|---|---|---|---|---|---|---|---|---|---|---|---|---|---|---|---|---|---|---|
| | $A_1$ | $T_2$ | $T_3$ | $C_4$ | $T_5$ | $A_6$ | $C_7$ | $C_8$ | $A_9$ | $G_{10}$ | $A_{11}$ | $G_{12}$ | $G_{13}$ | $T_{14}$ | $A_{15}$ | $C_{16}$ | $A_{17}$ | $A_{18}$ | $G_{19}$ | $G_{20}$ | A | G | G |
| G | 0.3 | 0.1 | 0.1 | 0.2 | 0.3 | 1.3 | 0.3 | 0.4 | 0.2 | 98.4 | 0.3 | 98.4 | 97.6 | 0.1 | 1.0 | 0.9 | 0.8 | 0.6 | 96.4 | 97.9 | 0.4 | 97.4 | 98.5 |
| A | 99.2 | 0.1 | 0.2 | 0.2 | 0.1 | 97.9 | 0.8 | 0.5 | 99.3 | 0.4 | 99.3 | 0.9 | 1.3 | 0.1 | 98.6 | 1.4 | 98.7 | 97.4 | 3.2 | 1.4 | 99.2 | 1.9 | 0.9 |
| T | 0.3 | 99.7 | 99.4 | 0.5 | 98.8 | 0.2 | 3.6 | 0.4 | 0.3 | 0.2 | 0.2 | 0.6 | 0.6 | 99.0 | 0.2 | 1.9 | 0.3 | 1.7 | 0.3 | 0.2 | 0.2 | 0.5 | 0.3 |
| C | 0.1 | 0.2 | 0.2 | 99.0 | 0.8 | 0.5 | 95.1 | 98.6 | 0.1 | 0.9 | 0.1 | 0.5 | 0.4 | 0.7 | 0.1 | 96.2 | 0.3 | 0.3 | 0.0 | 0.5 | 0.0 | 0.2 | 0.3 |

dABE

| | | | | | | | | | | | | | | | | | | | | | PAM | |
|---|---|---|---|---|---|---|---|---|---|---|---|---|---|---|---|---|---|---|---|---|---|---|
| | $A_1$ | $T_2$ | $T_3$ | $C_4$ | $T_5$ | $A_6$ | $C_7$ | $C_8$ | $A_9$ | $G_{10}$ | $A_{11}$ | $G_{12}$ | $G_{13}$ | $T_{14}$ | $A_{15}$ | $C_{16}$ | $A_{17}$ | $A_{18}$ | $G_{19}$ | $G_{20}$ | A | G | G |
| G | 0.4 | 0.1 | 0.1 | 0.2 | 0.1 | 49.6 | 0.4 | 0.4 | 2.3 | 98.4 | 0.3 | 98.6 | 97.9 | 0.1 | 0.8 | 0.7 | 0.7 | 0.5 | 96.9 | 97.9 | 0.3 | 97.8 | 98.7 |
| A | 99.3 | 0.1 | 0.2 | 0.1 | 0.1 | 49.7 | 0.7 | 0.4 | 97.3 | 0.5 | 99.2 | 0.6 | 1.2 | 0.0 | 98.7 | 1.1 | 98.9 | 97.9 | 2.8 | 1.2 | 99.5 | 1.7 | 0.7 |
| T | 0.2 | 99.7 | 99.4 | 0.5 | 99.2 | 0.2 | 3.3 | 0.4 | 0.3 | 0.1 | 0.3 | 0.6 | 0.6 | 99.3 | 0.4 | 1.9 | 0.1 | 1.5 | 0.3 | 0.4 | 0.2 | 0.3 | 0.3 |
| C | 0.1 | 0.2 | 0.3 | 99.0 | 0.6 | 0.5 | 95.5 | 98.8 | 0.1 | 0.9 | 0.2 | 0.5 | 0.4 | 0.5 | 0.1 | 96.4 | 0.2 | 0.1 | 0.0 | 0.5 | 0.0 | 0.2 | 0.3 |

**Figure S11 | (A)** Base editing window comparing ABE vs CBE and nCas9-BE vs dCas9-BE in the top edited live single cell isolated stable cell line.

A)

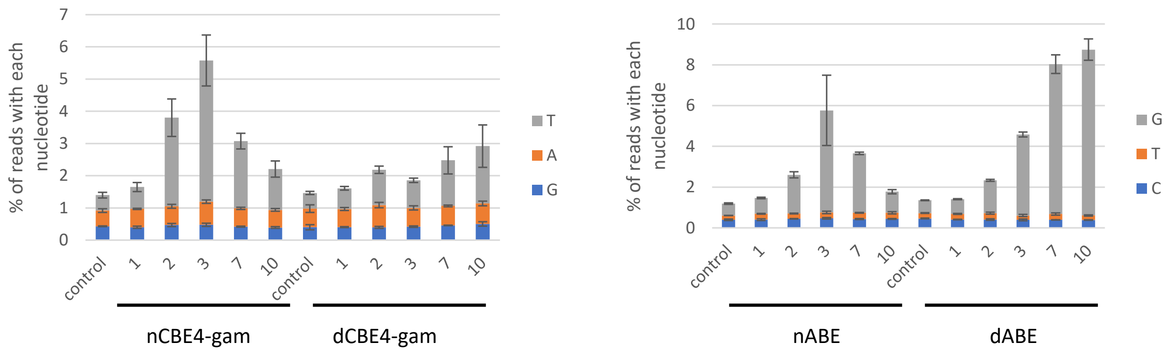


B)

HL1gR4 population control

| | $A_1$ | $T_2$ | $T_3$ | $C_4$ | $T_5$ | $A_6$ | $C_7$ | $C_8$ | $A_9$ | $G_{10}$ | $A_{11}$ | $G_{12}$ | $G_{13}$ | $T_{14}$ | $A_{15}$ | $C_{16}$ | $A_{17}$ | $A_{18}$ | $G_{19}$ | $G_{20}$ | A (PAM) | G (PAM) | G (PAM) | Indel % |
|---|---|---|---|---|---|---|---|---|---|---|---|---|---|---|---|---|---|---|---|---|---|---|---|---|
| G | 0.2 | 0.1 | 0.1 | 0.3 | 0.1 | 0.6 | 0.4 | 0.3 | 0.3 | 98.3 | 0.2 | 98.2 | 97.8 | 0.0 | 1.1 | 1.0 | 0.8 | 0.4 | 95.9 | 97.7 | 0.4 | 97.6 | 98.7 | 1.22 |
| A | 99.5 | 0.1 | 0.3 | 0.1 | 0.1 | 98.7 | 0.9 | 0.7 | 99.3 | 0.5 | 99.2 | 1.1 | 1.2 | 0.0 | 98.7 | 1.4 | 98.8 | 98.1 | 3.8 | 1.6 | 99.4 | 1.9 | 0.9 | |
| T | 0.2 | 99.7 | 99.4 | 0.4 | 99.3 | 0.2 | 3.6 | 0.5 | 0.3 | 0.2 | 0.3 | 0.3 | 0.6 | 99.2 | 0.1 | 1.8 | 0.2 | 1.2 | 0.2 | 0.3 | 0.1 | 0.5 | 0.3 | |
| C | 0.0 | 0.2 | 0.2 | 99.2 | 0.5 | 0.4 | 95.0 | 98.5 | 0.1 | 1.0 | 0.2 | 0.5 | 0.5 | 0.7 | 0.1 | 95.9 | 0.2 | 0.3 | 0.1 | 0.4 | 0.1 | 0.2 | 0.2 | |

nCBE4-gam

| | $A_1$ | $T_2$ | $T_3$ | $C_4$ | $T_5$ | $A_6$ | $C_7$ | $C_8$ | $A_9$ | $G_{10}$ | $A_{11}$ | $G_{12}$ | $G_{13}$ | $T_{14}$ | $A_{15}$ | $C_{16}$ | $A_{17}$ | $A_{18}$ | $G_{19}$ | $G_{20}$ | A (PAM) | G (PAM) | G (PAM) | Indel % |
|---|---|---|---|---|---|---|---|---|---|---|---|---|---|---|---|---|---|---|---|---|---|---|---|---|
| G | 0.2 | 0.0 | 0.2 | 0.4 | 0.1 | 0.5 | 0.4 | 0.4 | 0.4 | 98.1 | 0.3 | 98.4 | 97.6 | 0.2 | 0.9 | 1.0 | 0.7 | 0.5 | 95.1 | 98.0 | 0.4 | 97.5 | 98.4 | 1.41 |
| A | 99.4 | 0.2 | 0.2 | 0.4 | 0.1 | 98.9 | 0.9 | 0.6 | 99.2 | 0.6 | 99.1 | 0.9 | 1.3 | 0.1 | 98.8 | 1.2 | 98.8 | 98.1 | 4.4 | 1.4 | 99.5 | 2.1 | 1.0 | |
| T | 0.3 | 99.6 | 99.3 | 2.9 | 99.1 | 0.1 | 5.5 | 2.5 | 0.3 | 0.2 | 0.3 | 0.3 | 0.6 | 99.2 | 0.2 | 1.6 | 0.1 | 1.1 | 0.3 | 0.3 | 0.1 | 0.3 | 0.2 | |
| C | 0.1 | 0.2 | 0.3 | 96.2 | 0.7 | 0.4 | 93.0 | 96.5 | 0.1 | 1.0 | 0.2 | 0.3 | 0.5 | 0.5 | 0.1 | 96.3 | 0.3 | 0.3 | 0.1 | 0.4 | 0.1 | 0.2 | 0.4 | |

dCBE4-gam

| | $A_1$ | $T_2$ | $T_3$ | $C_4$ | $T_5$ | $A_6$ | $C_7$ | $C_8$ | $A_9$ | $G_{10}$ | $A_{11}$ | $G_{12}$ | $G_{13}$ | $T_{14}$ | $A_{15}$ | $C_{16}$ | $A_{17}$ | $A_{18}$ | $G_{19}$ | $G_{20}$ | A (PAM) | G (PAM) | G (PAM) | Indel % |
|---|---|---|---|---|---|---|---|---|---|---|---|---|---|---|---|---|---|---|---|---|---|---|---|---|
| G | 0.3 | 0.1 | 0.2 | 0.4 | 0.1 | 0.7 | 0.5 | 0.4 | 0.3 | 97.8 | 0.3 | 96.8 | 96.7 | 0.2 | 1.1 | 1.0 | 0.9 | 0.5 | 91.5 | 97.0 | 0.5 | 95.9 | 98.0 | 1.43 |
| A | 99.2 | 0.1 | 0.3 | 0.3 | 0.2 | 98.6 | 0.9 | 0.6 | 99.3 | 0.8 | 99.1 | 1.2 | 1.8 | 0.1 | 98.4 | 1.5 | 98.6 | 98.0 | 8.2 | 1.8 | 99.3 | 3.5 | 1.4 | |
| T | 0.3 | 99.5 | 99.2 | 1.7 | 99.0 | 0.2 | 4.6 | 1.5 | 0.3 | 0.2 | 0.4 | 1.4 | 1.0 | 98.9 | 0.4 | 2.3 | 0.2 | 1.2 | 0.3 | 0.4 | 0.2 | 0.4 | 0.3 | |
| C | 0.1 | 0.4 | 0.3 | 97.6 | 0.7 | 0.5 | 93.9 | 97.4 | 0.1 | 1.1 | 0.1 | 0.6 | 0.5 | 0.8 | 0.1 | 95.2 | 0.3 | 0.2 | 0.1 | 0.9 | 0.1 | 0.2 | 0.3 | |

nABE

| | $A_1$ | $T_2$ | $T_3$ | $C_4$ | $T_5$ | $A_6$ | $C_7$ | $C_8$ | $A_9$ | $G_{10}$ | $A_{11}$ | $G_{12}$ | $G_{13}$ | $T_{14}$ | $A_{15}$ | $C_{16}$ | $A_{17}$ | $A_{18}$ | $G_{19}$ | $G_{20}$ | A (PAM) | G (PAM) | G (PAM) | Indel % |
|---|---|---|---|---|---|---|---|---|---|---|---|---|---|---|---|---|---|---|---|---|---|---|---|---|
| G | 0.5 | 0.1 | 0.2 | 0.5 | 0.1 | 3.0 | 0.4 | 0.4 | 0.5 | 97.9 | 0.3 | 97.3 | 96.8 | 0.1 | 1.0 | 1.2 | 1.0 | 0.6 | 90.1 | 96.8 | 0.6 | 96.7 | 97.9 | 1.49 |
| A | 99.1 | 0.3 | 0.2 | 0.4 | 0.2 | 96.2 | 0.9 | 0.6 | 98.9 | 0.9 | 98.9 | 1.4 | 2.0 | 0.2 | 98.5 | 1.5 | 98.6 | 98.0 | 9.5 | 2.1 | 99.1 | 2.7 | 1.5 | |
| T | 0.3 | 99.4 | 99.3 | 0.7 | 98.7 | 0.3 | 3.4 | 0.7 | 0.4 | 0.2 | 0.4 | 0.7 | 0.8 | 98.5 | 0.4 | 2.2 | 0.3 | 1.1 | 0.4 | 0.4 | 0.2 | 0.4 | 0.3 | |
| C | 0.1 | 0.4 | 0.3 | 98.3 | 0.9 | 0.5 | 95.1 | 98.3 | 0.1 | 1.1 | 0.2 | 0.5 | 0.5 | 1.2 | 0.1 | 95.3 | 0.2 | 0.3 | 0.1 | 0.7 | 0.1 | 0.2 | 0.4 | |

dABE

| | $A_1$ | $T_2$ | $T_3$ | $C_4$ | $T_5$ | $A_6$ | $C_7$ | $C_8$ | $A_9$ | $G_{10}$ | $A_{11}$ | $G_{12}$ | $G_{13}$ | $T_{14}$ | $A_{15}$ | $C_{16}$ | $A_{17}$ | $A_{18}$ | $G_{19}$ | $G_{20}$ | A (PAM) | G (PAM) | G (PAM) | Indel % |
|---|---|---|---|---|---|---|---|---|---|---|---|---|---|---|---|---|---|---|---|---|---|---|---|---|
| G | 0.4 | 0.1 | 0.2 | 0.4 | 0.1 | 8.2 | 0.5 | 0.5 | 0.9 | 97.7 | 0.3 | 96.8 | 96.3 | 0.1 | 1.0 | 1.2 | 0.9 | 0.5 | 89.0 | 96.8 | 0.4 | 96.2 | 97.7 | 1.41 |
| A | 99.2 | 0.3 | 0.3 | 0.3 | 0.2 | 91.1 | 0.9 | 0.5 | 98.6 | 0.8 | 99.0 | 1.5 | 2.0 | 0.1 | 98.5 | 1.6 | 98.7 | 98.2 | 10.7 | 1.8 | 99.3 | 3.2 | 1.5 | |
| T | 0.3 | 99.4 | 99.1 | 0.8 | 98.8 | 0.3 | 3.4 | 0.7 | 0.3 | 0.2 | 0.4 | 1.1 | 1.0 | 98.8 | 0.4 | 2.3 | 0.3 | 1.0 | 0.4 | 0.3 | 0.2 | 0.4 | 0.3 | |
| C | 0.1 | 0.4 | 0.3 | 98.4 | 0.9 | 0.4 | 95.1 | 98.3 | 0.1 | 1.2 | 0.2 | 0.6 | 0.6 | 1.0 | 0.1 | 95.0 | 0.2 | 0.3 | 0.1 | 1.0 | 0.1 | 0.2 | 0.4 | |

**Figure S12 | Base editing purity in HEK 293T targeting LINE-1. (A)** Purity of deamination at target nucleotide (left) CBE, (right) ABE. **(B)** Base editing activity across gRNA target sequence at day seven.

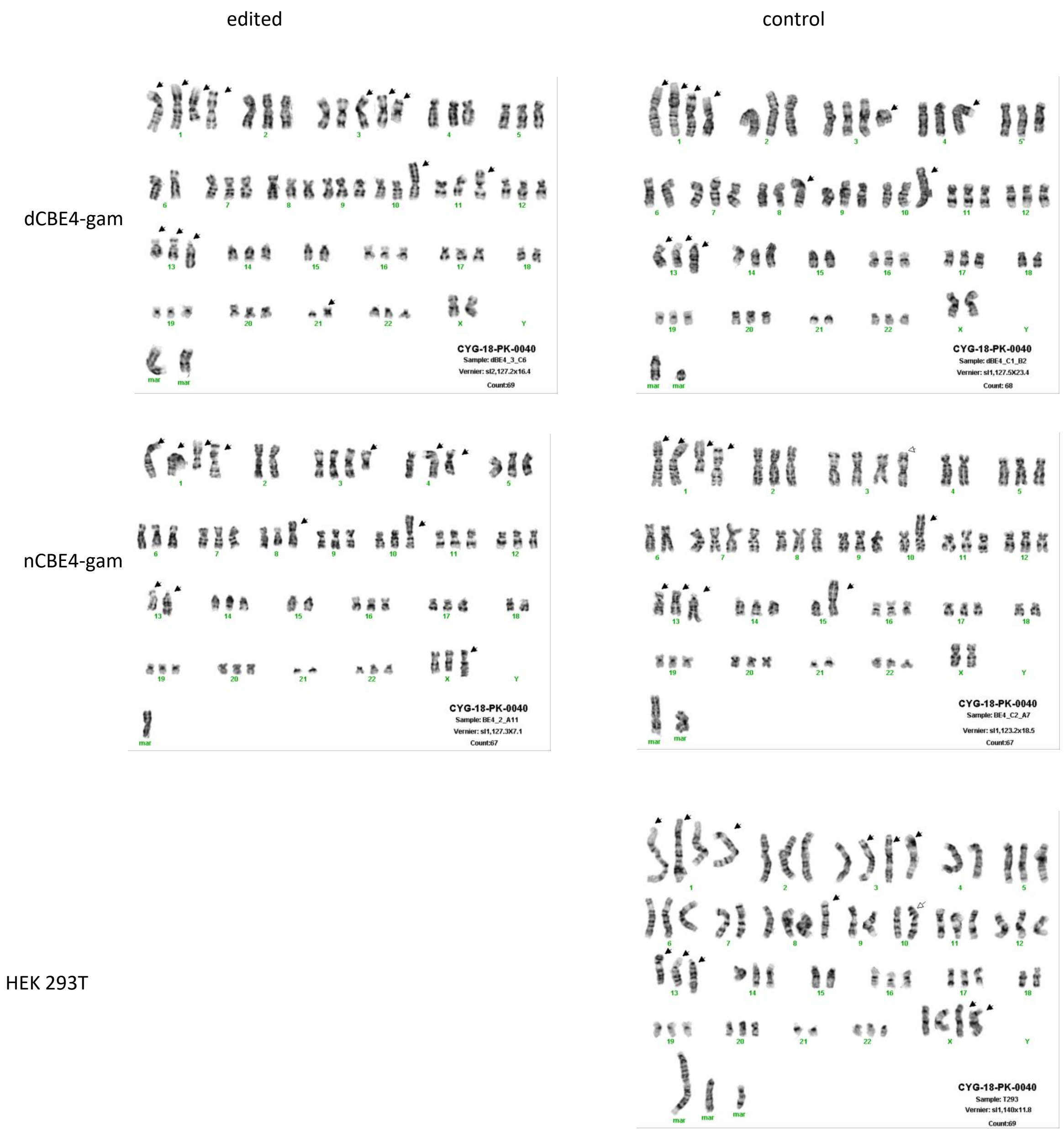


**Figure S13 | Karyotype analysis after CBE4-gam editing. (A)** Karyotype chromosome presentations for each of the genome edited clones and controls.

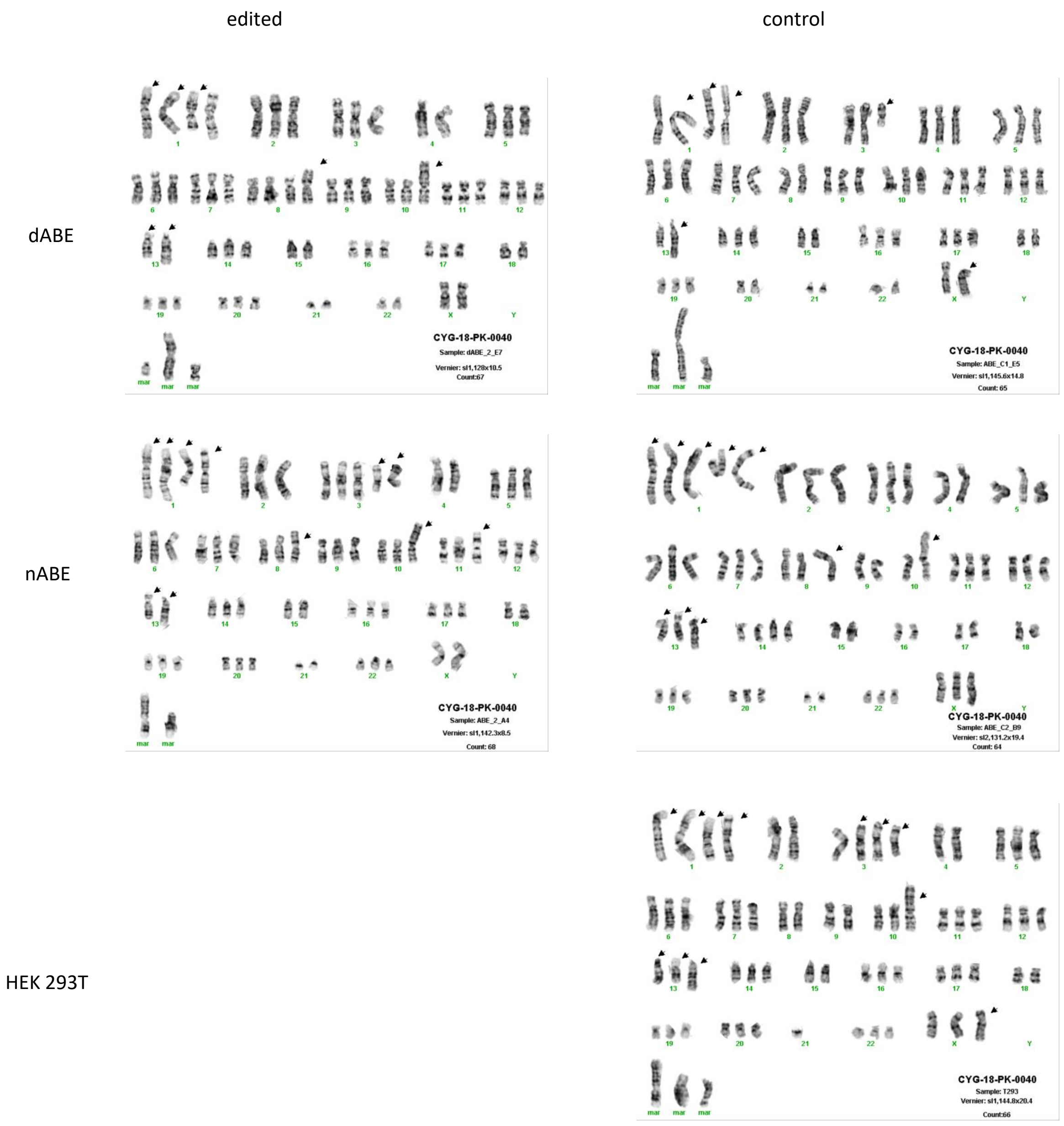


**Figure S14 | Karyotype analysis after ABE editing. (A)** Karyotype chromosome presentations for each of the genome edited clones and controls.

**A)**

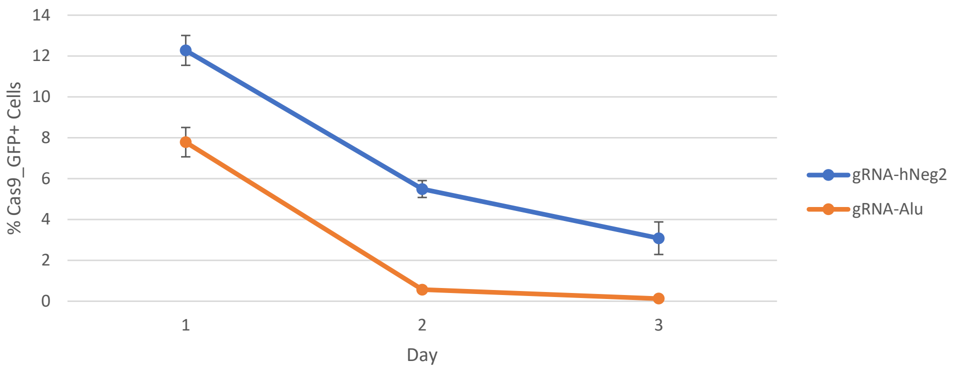


**Figure S15 | TE gRNAs are highly toxic in human iPSCs (A)** Percentage Cas9_GFP+ cells over time after transfection with TE or control gRNAs.

A)

wt

| | $T_3$ | $C_4$ | $T_5$ | $A_6$ | $C_7$ | $C_8$ | $A_9$ |
|---|---|---|---|---|---|---|---|
| G | 0.00 | 0.00 | 0.00 | 0.80 | 0.40 | 0.40 | 0.40 |
| A | 0.00 | 0.80 | 0.00 | 98.80 | 0.40 | 0.40 | 99.20 |
| T | 99.20 | 1.60 | 98.80 | 0.00 | 2.80 | 0.40 | 0.00 |
| C | 0.80 | 97.60 | 0.80 | 0.40 | 96.40 | 98.90 | 0.40 |

cK

| | $T_3$ | $C_4$ | $T_5$ | $A_6$ | $C_7$ | $C_8$ | $A_9$ |
|---|---|---|---|---|---|---|---|
| G | 0.3 | 0.3 | 0.0 | 1.5 | 0.3 | 0.6 | 0.7 |
| A | 0.0 | 0.0 | 0.0 | 97.4 | 0.7 | 0.7 | 99.3 |
| T | 99.4 | 2.1 | 97.8 | 0.3 | 5.1 | 2.7 | 0.0 |
| C | 0.3 | 97.6 | 2.2 | 0.8 | 94.0 | 96.0 | 0.0 |

cKA5

| | $T_3$ | $C_4$ | $T_5$ | $A_6$ | $C_7$ | $C_8$ | $A_9$ |
|---|---|---|---|---|---|---|---|
| G | 0.6 | 0.3 | 0.0 | 2.2 | 1.0 | 1.3 | 1.1 |
| A | 0.0 | 0.0 | 0.0 | 97.1 | 1.0 | 1.0 | 98.9 |
| T | 98.7 | 1.3 | 98.6 | 0.6 | 4.1 | 1.9 | 0.0 |
| C | 0.6 | 98.4 | 1.4 | 0.0 | 93.8 | 95.7 | 0.0 |

cKD5

| | $T_3$ | $C_4$ | $T_5$ | $A_6$ | $C_7$ | $C_8$ | $A_9$ |
|---|---|---|---|---|---|---|---|
| G | 0.7 | 0.0 | 0.0 | 0.3 | 0.7 | 0.4 | 0.4 |
| A | 0.4 | 1.1 | 0.0 | 99.3 | 0.4 | 0.7 | 99.2 |
| T | 98.6 | 3.6 | 99.6 | 0.0 | 4.7 | 3.7 | 0.0 |
| C | 0.3 | 95.3 | 0.4 | 0.3 | 93.9 | 95.2 | 0.4 |

B)

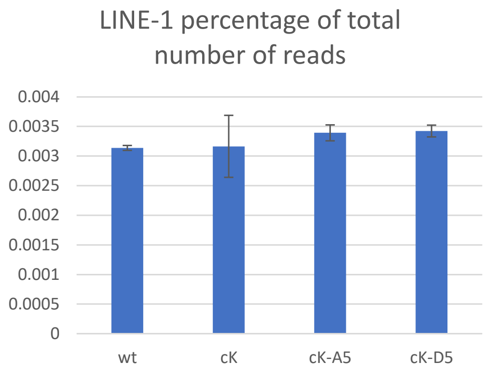


C)

| | clone K vs. wt | cKA5 vs. wt | cKD5 vs. wt |
|---|---|---|---|
| downregulated genes | 3055 | 2721 | 3012 |
| not significant | 7442 | 8114 | 7555 |
| upregulated genes | 2913 | 2591 | 2848 |

D)

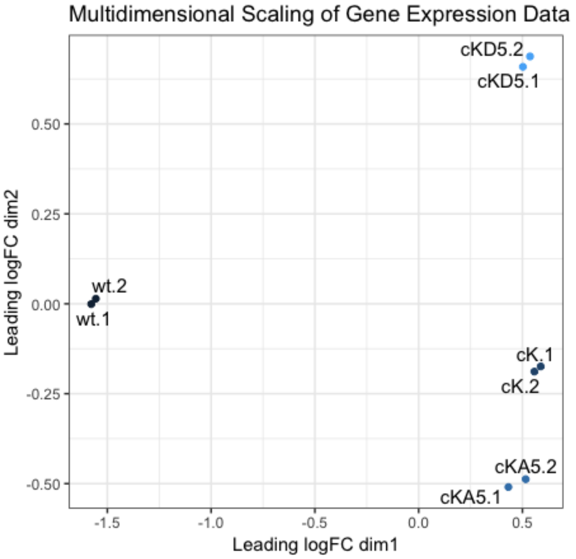


**Figure S16 | LINE-1 RNA expression in KO clones (A)** Base editing activity detected in RNA transcripts of clone K (cK), clone K-A5 (cKA) and clone K-D4 (cKD5) within the gRNA target sequence **(B)** Percentage of LINE-1 reads relative to total number of reads. Error bars represent standard deviation between biological duplicates. **(C)** Summary of differentially expressed genes as determined by the exact test. **(D)** Multidimensional scaling plot where distance corresponds to leading log-fold count changes between the RNA samples.

### SUPPLEMENTAL TABLES

| | C-deaminase | A-deaminase | UGI | Nick | Mu gam |
|---|---|---|---|---|---|
| dCBE1[1] | X | | | | |
| dCBE2[1] | X | | X | | |
| nCBE3[1] | X | | X | X | |
| nCBE4[2] | X | | X2 | X | |
| nCBE4-gam[2] | X | | X2 | X | X |
| dCBE4* | X | | X2 | | |
| dCBE4-gam* | X | | X2 | | X |
| nABE[3] | | X | | X | |
| dABE* | | X | | | |

*Synthesized and tested in this study

[1]Komor et al. (2016) *Nature* **533**(7603):420-4

[2]Komor et al. (2017) *Sci Adv* **3**(8):eaao4774

[3]Guadelli et al. (2017) *Nature* **551**:464-71

Table S1: Evolution of base editors

| Manuscript name | Addgene name | Addgene # |
|---|---|---|
| pSB700 | pSB700 | 64046 |
| pSB700_mCherry | - | - |
| pSB700_Puro | - | - |
| SaCas9_gRNA | BPK2660 | 70709 |
| pCas9_GFP | pCas9_GFP | 44719 |
| hCas9 | hCas9 | 41815 |
| nCBE2 | pCMV_BE2 | 73020 |
| nCBE3 | pCMV_BE3 | 73021 |
| nCBE4 | BE4 | 100802 |
| nCBE4-gam | BE4-gam | 100806 |
| nABE | pCMV_ABE7.10 | 102919 |
| SaCas9 | pX600-AAV-CMV::NLS-SaCas9-NLS-3xHA-bGHpA | 61592 |
| Sa-nCBE4-gam | SaBE4-gam | 100809 |
| SaKKH-nCBE4 | pJL-SaKKH-BE3 | 85170 |
| dCBE4 | - | |
| dCBE4-gam | dCBE4-gam | |
| dABE | dABE | |

**Table S2:** List of DNA editors

| Name | gRNA | PAM | Cas9 species |
|---|---|---|---|
| Non human | GAGACGATTAATGCGTCTCG | NGG | SpCas9 |
| S1 | GATGACAGGCAGGGGCACCG | CGG | SpCas9 |
| HL1 gR1 | AACGAGACAGAAAGTCAACA | AGG | SpCas9 |
| HL1 gR2 | TCAGTTTCCATGTAGTTGAG | CGG | SpCas9 |
| HL1 gR3 | TATGTACCCAGTAGTCATTC | AGG | SpCas9 |
| HL1 gR4 | ATTCTACCAGAGGTACAAGG | AGG | SpCas9 |
| HL1 gR5 | TTGAACCAGCCTTGCATCCC | AGG | SpCas9 |
| HL1 gR6 | GGGTATTCAATTAGGAAAAG | AGG | SpCas9 |
| EN gR1 | GACTCCCACACATTAATAAT | GGG | SpCas9 |
| HL1 gR46 | GCTTAGGTAAACAAAGCAGC | | SpCas9 |
| EN gR9 | ATTTTGGAATAGGTGTGGTG | TGG | SpCas9 |
| RT gR1 | ATTCAGTATGATATTGGCTG | TGG | SpCas9 |
| RT gR3 | CCTAGGAATCCAACTTACAA | GGG | SpCas9 |
| Z8 gR2 | AAAAAGAGTCCAGGACCAGA | TGG | SpCas9 |
| Alu | CAGGCGTGAGCCACCGCGCC | CGG | SpCas9 |
| Sa Non human | GAGACGATTAATGCGTCTCG | NGG | SaCas9 |
| HERV env11 | GAGGCACATCCAACAGTTAG | TAGGG | SaCas9 |

**Table S3:** gRNAs used in this study

ILMN – F: 5'- CTTTCCCTACACGACGCTCTTCCGATCT -3'
ILMN – R: 5'- GGAGTTCAGACGTGTGCTCTTCCGATCT -3'

| gRNA Name | PrimerF | PrimerR |
|---|---|---|
| S1 | TCAAGATGGCTGACAAAG | GCACCAGAGTCTCCGCTTTA |
| HL1 gR1 | AGACTCCCACACATTAATAATGGG | TGATTTGGGGTGGAGAGTTCTG |
| HL1 gR2 | AGTGCAATCAAACTAGAACTCAGG | CCCTCTACACACTGCTTTGAATG |
| HL1 gR3 | AGTGCAATCAAACTAGAACTCAGG | CCCTCTACACACTGCTTTGAATG |
| HL1 gR4 | AAGAGTCCAGGACCAGATGGAT | CCCGGCTTTGGTATCAGAATG |
| HL1 gR5 | CTTATCCACCATGATCAAGTGGG | CTGCATCTATTGAGATAATCATGTGG |
| HL1 gR6 | GTTCTGGCCAGGGCAATCAG | CCTGAGACTTTGCTGAAGTTGC |
| HL1 gR46 | AACTGCAAGGCGGCAACGAG | AGAGGTGGAGCCTACAGAGG |
| EN gR1 | CCAATACAGGAGCACCCAGATT | TGATTTGGGGTGGAGAGTTC |
| EN gR9 | CAGAACTCTCCACCCCAAAT | CCTGAGTTCTAGTTTGATTG |
| RT gR1 | CCACATGATTATCTCAATAG | GAGGGCATCCCTGTCTTGTG |
| RT gR3 | GCAACTTCAGCAAAGTCTCA | GTAGTTCTCCTTGAAGAGGTCC |
| EN (dual gRNA) | CCAATACAGGAGCACCCAGATT | CCCTCTACACACTGCTTTGAATG |
| RT (dual gRNA) | CCACATGATTATCTCAATAG | GTAGTTCTCCTTGAAGAGGTCC |
| ENRT (dual gRNA) | CAGAACTCTCCACCCCAAATC | CCCGGCTTTGGTATCAGAATG |
| shEN (dual gRNA) | CAGAACTCTCCACCCCAAATC | CCCTCTACACACTGCTTTGAATG |
| HERV env11 | AATACCACCCTCACTGGGCT | CAGATTGGAAACAAGAGGTCC |

**Table S4:** NGS primers list

| | BE4_2_A11 | BE4_C2_A7 | dBE4_3_C6 | dBE4_C1_B2 | ABE_2_A4 | ABE_C2_B9 | dABE_2_E7 | dABE_C1_E2 | 293T9(CYG-18-PK-0040) |
|---|---|---|---|---|---|---|---|---|---|
| -X | | | | x | x | | x | x | |
| add(X)(q28) | x | x | x | | x | x | | | x |
| der(X)add(X)(p11.2)add(X)(q28) | | | | | | | | x | |
| add(1)(p36.1) | x | x | x | x | x | x | x | | x |
| add(1)(q42) | xx | xx | xx | xx | xx | xx | xx | | xx |
| del(1)(q31) | x | x | x | x | | x | x | | x |
| i(1)(p10) | | | | | | | | x | |
| add(1)(q21) | | | | | | | | x | |
| -2 | | | | | | | | | |
| add(3)(p13) | | | | | | | | x | |
| add(3)(p24) | | | xx | | | | | | x |
| del(3)(p22) | x | | | | x | | | | |
| add(3)(q12) | | | x | x | | | | | x |
| del(3)(q22) | x | | | | x | x | | x | |
| add(4)(p15) | x | | | | | | | | |
| del(4)(q31) | x | | | | | | | | |
| -4 | | x | | | x | x | x | | x |
| add(8)(p21) | x | x | x | x | x | x | x | | x |
| -9 | | | | | | x | | | |
| add(10)(p11) | | | | | | | | | |
| add(10)(p13) | x | x | x | x | x | x | x | | x |
| add(11)(p15) | | | x | | x | | | | |
| add(13)(p11) | xx | xx | xx | xx | xx | xx | xx | | xx |
| add(13)(q34) | x | x | x | x | x | x | x | | x |
| -13 | | | | | | | | | |
| add(14)(p11.2) | | | x | x | | | | x | |
| -15 | x | x | x | x | x | x | x | | x |
| add(15)(p11.2) | | x | | | | | | | |
| -18 | x | x | x | x | x | x | x | | x |
| -21 | x | x | | x | x | x | x | | x |
| -22 | | | | | | | x | | |
| i(21)(q10) | | | x | | | | | | |
| mar | x-xx | x-xx | x-xxx | x-xx | xx-xxx | x-xx | x-xxx | x-xxx | x-xxxx |

### Table S5: Karyotype chromosomal abnormality list

#### SUPPLEMENTARY RESULTS

##### Single cell analysis of LINE-1 dual gRNA disrupted cells

The PCR amplicons of dual gRNA combinations from the previous experiments were too large to include both the mutated and full-length bands together for Illumina NGS. To overcome this, a shorter pair of LINE-1 targeting gRNAs, called short EN (shEN), was used that permits both regions to be sequenced together. 293Ts were transfected with pCas9_GFP and the shEN gRNA pair (ENgR9 and HL1gR3) in the pSB700mCherry gRNA expression vector (*fig. S3A*). GFP and mCherry double-positive single cells were FACS-sorted into gDNA extraction solution. 303 individual cells were screened after FACS sorting and LINE-1 NGS analysis. Of those wells with an amplicon, 83.24% had a visually detectable deletion band with a range of intensities from barely observable to stronger than the wild type non-mutated band (*fig. S3B*). Bulk-transfected cells had a dual gRNA deletion frequency of 2.7%, the FACS-enriched double-positive cell population was edited at 11.19%, and the mean editing of single-cell-derived amplicons was ~50.17% (*fig. S3C*). The editing frequency appears to be bimodal as previously reported in the set of PERV editing papers, with experiments first in transformed cells(*8*) – achieving 62 indels – and then later in healthy born piglets(*9*) – with all 25 PERVs knocked-out. At first it seems contradictory that the population bulk gDNA editing efficiency is 11.19% and the single cell average is 50.1% but this assumes that each single cell had a full nuclear genome. The highest edited samples most likely already had degraded their genomes due to the thousands of concurrent cuts to every chromosome received; thus, each single cell observed was contributing unequally to the bulk population.

##### nBE activity confirmed at LINE-1 for both nABE and nCBE

We tested the efficiency of deamination at LINE-1 using base editors. We designed and tested LINE-1 targeting gRNAs (HL1gR1-6 [*table S1*]) that generate a STOP codon early in ORF-2 using C->T deamination. HEK 293T cells were transfected with nCBE3 and gRNAs individually. Deamination events were detected at each of the six gRNA target loci above the background levels of editing observed (~0.05% – 0.67%) in mock transfected populations of cells (*fig. S4A*). These same CBE gRNAs were also compatible with ABEs as they contain at least one adenine within their target window. Base editing with nCBE4-gam and nABE was detected in the population in 4/5 gRNAs for CBE (*fig. S4B*) and 4/5 gRNAs for ABE (*fig. S4C*). nABE had the highest editing efficiency using HL1gR6 at 4.94% or ~1290 loci genome wide. HL1gR4 was chosen as the best target for future studies as its signal to background error ratio was the lowest of all the LINE-1 amplicons/gRNAs tested, and the HL1gR4 was among the most efficient. The HL1gR4 target also contains three C's within its target window that are all efficiently co-edited as a clear watermark of mutation. An Alu-targeting gRNA resulted in increased cell survival when using nCBE3 compared to Cas9 (*fig. S5*).

##### RNA-seq in LINE-1 KO clones

We did not observe downregulation of LINE-1 RNA expression levels in edited clones. In *Fig. S16B* the number of RNA reads obtained through the standard deamination analysis pipeline, averaged over the 20 nt protospacer sequence and normalized the read counts by dividing by the size of their respective libraries, are displayed. A list of predicted differentially expressed genes in the edited clones compared to the wild type is found in supplementary data S1, and numbers of up and down regulated genes is found in figure S16C. Multidimensional scaling of the gene expression data (FigS16D), where the distance between the samples corresponds to leading log-fold-changes between the RNA samples, shows a clear separation between the wild type and the three edited samples. Since the wild type control samples, however, did not undergo a comparable procedure of transfection and cell sorting, we cannot conclude that the observed differences in gene expression are due to LINE-1 editing.

#### DATA AND MATERIALS AVAILABILITY

Key plasmids developed during this study have been submitted to Addgene: pSB700_HL1 gR4 (# 124450), dABE (# 124447) and dCBE4-gam (# 124449). All NGS data used for the figures and supplementary figures have been made available at SRA BioProject Accession #PRJNA515875 and #PRJNA518077 for 293T and PGP1 respectively.
